# IL-17RC signaling connects intestinal microbiota and neuroimmune interactions in atherosclerosis

**DOI:** 10.64898/2026.03.06.710205

**Authors:** Aleksandra M. Mazitova, Jiani Zhu, Richard Rodrigues, Kaitlyn Nguyen, Meghan Terrell, Pavithra Nedumaran, Jacob Alltucker, Mingtian Che, Kelsey E. Jarrett, Kevin P Downs, Christian Stehlik, Thomas Q. de Aguiar Vallim, Andrew Kossenkov, Bernd Schnabl, Simon Knott, Giorgio Trinchieri, Amiran Dzutsev, Sergei I. Grivennikov, Ekaterina K. Koltsova

## Abstract

While dysbiosis and inflammation were previously implicated in cardiovascular diseases, the circuits of how microbiota drives distant perivascular innervation, neuroinflammation and atherosclerosis remains unknown. Here, we report that IL-17RC signaling in intestine protects from atherosclerosis controlling intestinal barrier and microbiota, and loss of IL-17RC in intestinal epithelial cells alters microbiota, enhances perivascular innervation and aortic inflammation, augmenting the disease. Neuronal outgrowth is functionally dependent on microbiota and is essential for neuroinflammation and augmentation of atherosclerosis as chemical denervation reduces inflammation, macrophage activation and disease progression. Microbiota-dependent IL-17A producing γδ T cells accumulate in aorta to promote neuronal outgrowth and activation that can be reversed by γδ T cell blockade. Perivascular neuron activation is further dependent on cell autonomous IL-17 signaling as IL-17RC ablation in sympathetic neurons protected mice from microbiota-driven atherosclerosis. Together, our data illuminate how intestinal cytokine signaling distantly restrains neuroimmune interactions in aorta and uncovers a novel link between IL-17 signaling, microbiota, perivascular innervation and neuroimmune pro-inflammatory crosstalk instrumental for atherosclerosis progression.

**Summary:** IL-17RC signaling regulates intestinal dysbiosis and perivascular neuronal outgrowth that modulates inflammation in atherosclerosis.

## INTRODUCTION

Atherosclerosis is the most prevalent form of cardiovascular disease (CVD)^1^ where inflammation plays a key role^2^. Progressive growth of atherosclerotic plaques is invariably associated with increased local and systemic levels of pro-inflammatory cytokines that regulate the inflammatory milieu^3,4^ and represent key mediators of cell-to-cell communication in the aorta and growing plaque^5^. Atherosclerosis is further promoted by systemic factors such as poor diet and stress; however, how these mechanisms can potentially interact and intersect with the regulation of the immune microenvironment in the aorta remains elusive.

Interleukin IL-17A (IL-17) has well-established pro-inflammatory roles in multiple inflammatory diseases, including arthritis, psoriasis, inflammatory bowel disease (IBD), and cancer^6,7^. Elevated levels of IL-17A and accumulation of IL-17A producing CD4 T cells in atherosclerotic vessels have been reported in mice and humans^8,9^. However, both pro-atherogenic and atheroprotective effects of IL-17, even in the same models, were described upon genetic or pharmacological inhibition of IL-17A/IL-17RA signaling^8–16^. IL-17 receptors, including IL-17RA and IL-17RC, are expressed by immune, stromal, and epithelial cells indicating a possibility of distinct cell type-specific responses to IL-17. Cytokine signaling regulates intestinal epithelial cells (IEC) and epithelial barrier function^17,18^, but the impact of IEC-specific IL-17 cytokine signaling as a regulator of host-microbiota interactions in systemic distant effects on aortic inflammation and atherosclerosis have never been investigated.

Recent studies demonstrated a reciprocal connection between the peripheral nervous system and immune cells, particularly in the context of intestinal inflammation^19,20^ and cancer^21^. Neurons express cytokine receptors^19^, and IL-17 was suggested to regulate the function of neurons in the brain and skin^22–25^. However, role of cytokines in the regulation of neurons in the aorta and link between IL-17 signaling, microbiota, and peripheral neuroimmune interactions in atherosclerosis remain enigmatic.

Here we report that unexpectedly IL-17RC signaling in intestinal epithelium protects from diet-induced atherosclerosis, as ablation of IL-17A/F specific receptor IL-17RC in IEC significantly exacerbated the disease. IL-17 dependent barrier-induced changes in microbiota led to distal control of atherosclerosis via heightened accumulation of IL-17A producing γδ T cells in the aortas, that in turn regulated expansion and activation of peripheral sympathetic neurons. Excessive activation of these neuroimmune circuits led to augmentation of local inflammation, macrophage activation and lipid uptake, enhancing atherosclerosis development. These atherosclerosis-promoting neuroinflammatory interactions were diet-dependent, driven by microbiota and further reversible by blockade of γδ T cells, genetic ablation of IL-17RC in TH^+^ sympathetic neurons or pharmacological inhibition of neurons. Overall, our study establishes that diet- and inflammation-induced perturbations of intestinal barrier and microbiota lead to systemic and aorta-specific activation of “γδ T cell-IL-17A- neuron-inflammation” pathway which promotes a common form of cardiovascular disease. This pathway mechanistically links multiple risk factors for atherosclerosis including diet, microbes, inflammation and stress.

## RESULTS

### Intestinal epithelial cell-specific IL-17RC deficiency aggravates atherosclerosis

Heterodimeric IL-17 receptor complex consists of two subunits, IL-17RA, which is a shared for several other cytokines, and IL-17RC, which is restricted to IL-17A and IL-17F^26^. Conflicting and opposite results even in the same atherosclerosis models have been reported for the roles of IL-17A^8–16^. To determine how IL-17A/IL-17RC signaling in intestinal epithelial cells (IEC) is implicated in regulation of atherosclerosis, we generated *ll17rc^fl/fl^* mice (Figure S1A)^27^ and produced *Ldlr^−/−^ll17rc^fl/fl^VilCre^+^*(*Ldlr^−/−^ll17rc*^ΔIEC^) and Cre^−^ (*Ldlr^−/−^ll17rc*^WT^) littermate control mice on atherosclerosis-prone *Ldlr^−/−^* background. The efficiency of IL-17RC gene deletion in the intestine was confirmed by Q-RT-PCR (Figure S1B). *Ldlr^−/−^ll17rc*^ΔIEC^ and *Ldlr^−/−^ll17rc*^WT^ littermate mice were separately housed upon weaning and fed with Western Diet (WD) for 16 weeks to induce atherosclerosis. No difference in body weight, circulating lipids, blood pressure and heart rate had been detected (Figure S1C, D, E, F). Analysis of atherosclerotic lesions revealed that inactivation of IL-17RC signaling in intestinal epithelial cells significantly enhanced atherosclerosis in both male and female WD-fed *Ldlr^−/−^*mice, suggesting its atheroprotective role in the gut (Figure 1A, B). Transcriptomic analysis of aortas isolated from separately housed WD-fed *Ldlr^−/−^ll17rc*^ΔIEC^ and *Ldlr^−/−^ll17rc*^WT^ mice revealed upregulation of multiple inflammatory pathways including IL-17, IL-6 and myeloid cell activation (Figure 1C and D). Furthermore, we also surprisingly found strong upregulation of gene expression signature consistent with neuronal outgrowth and activation (Figure 1C and D). Particularly, expression of genes associated with synaptic transmission (*Cadm1, Arrb2*), axon neogenesis and differentiation (*Ngfr, Dclk1r, Nrp2, Slitrk1*), cholinergic receptors (*Chrnb4, Chrna3*), *Gap43* (regenerating neurons), adrenergic neurons (*Th)* and Norepinephrine synthesis (*Dbh, Ddc, Slc6a2*) were significantly elevated in aortas from *Ldlr^−/−^ll17rc*^ΔIEC^ mice (Figure 1C and D). The results were further validated by Q-RT-PCR in aortas of separate cohorts of mice (Figure 1E and F). No difference was found in neuronal gene expression in spleen (Figure S1G) and we detected the downregulation of selected neuronal genes in ileum of *Ldlr^−/−^ll17rc*^ΔIEC^ compared to *Ldlr^−/−^ll17rc*^WT^ mice (Figure S1H), suggesting aorta-specific effect.

**Figure 1.**
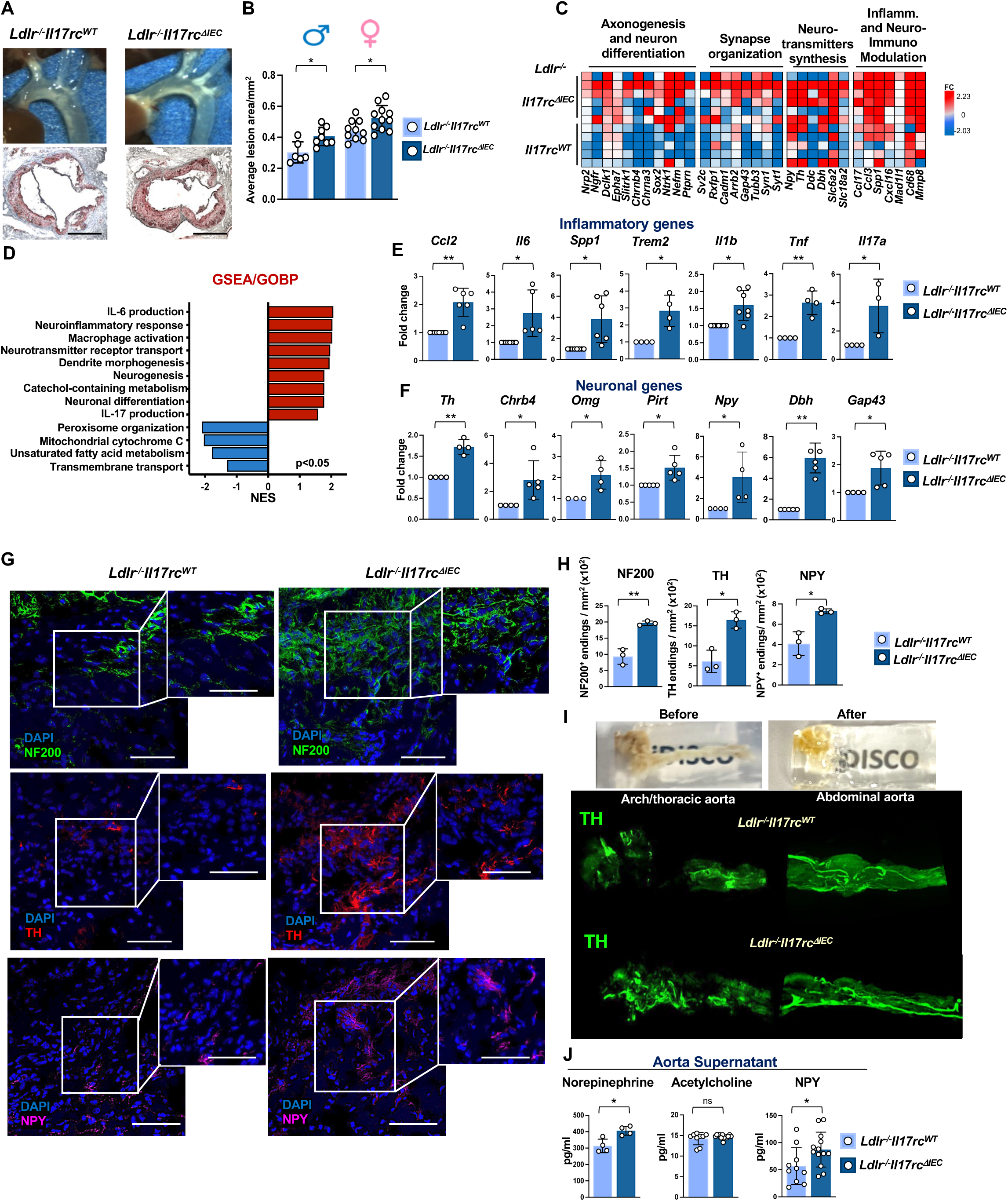
Epithelial cell IL-17RC protects from atherosclerosis and controls aortic inflammation and neuronal expansion. Separately housed *Ldlr^−/−^ll17rc^WT^* and *Ldlr^−/−^ll17rc*^Δ^*^IEC^* mice were allowed to develop atherosclerosis for 16 weeks of WD feeding. **A**. Images of aortic arch and representative images of aortic root sections. Scale bar is 1000 μm. **B**. Quantitative comparison of atherosclerotic lesion sizes in *Ldlr^−/−^ll17rc^WT^* and *Ldlr^−/−^ll17rc*^Δ^*^IEC^* mice (n = 6-11). **C**. Bulk RNA sequencing analysis of aortic tissues with a heatmap of differentially expressed genes. FC-Fold changes. **D** Gene Set Enrichment Analysis (GSEA) of GO Biological Processes (GOBP) from bulk RNA-seq of aortas from *Ldlr^−/−^ll17rc^WT^* and *Ldlr^−/−^ll17rc*^Δ^*^IEC^* mice. Red –pathways upregulated in *Ldlr^−/−^ll17rc*^Δ^*^IEC^* compared to *Ldlr^−/−^ll17rc^WT^*, Blue - pathways downregulated in *Ldlr^−/−^ll17rc*^Δ^*^IEC^* compared to *Ldlr^−/−^ll17rc^WT^.* Expression of selected Inflammatory **(E)** and neuronal **(F)** genes in aortas housed *Ldlr^−/−^ll17rc^WT^* (n =3-7) and *Ldlr^−/−^ll17rc*^Δ^*^IEC^* (n=3-7) mice. Gene expression was normalized to *Rpl32* and then to gene expression in *Ldlr^−/−^ll17rc^WT^.* Data are mean ± SEM from at least 3 independent experiments. *p<0.05, **p<0.01. Student’s t-test. Representative images (**G**) and quantification (**H**) of aortic root sections from *Ldlr^−/−^ll17rc^WT^* and *Ldlr^−/−^ll17rc*^Δ^*^IEC^* mice stained for neurons. Neurofilament 200 (NF200) (Green), Tyrosine Hydroxylase (TH) (Red) and Neuropeptide Y (NPY) (Magenta). Scale bar is 50 μm. Representative from 3 independent experiments. *p<0.05, **p<0.01. Student’s t-test**. I.** Representative iDISCO-cleared images of mouse aortas *Ldlr^−/−^ll17rc^WT^* and *Ldlr^−/−^ll17rc*^Δ^*^IEC^* mice **(top)** and Lightsheet microscopy imaging of iDISCO-cleared aortas stained for TH **(bottom). J.** Norepinephrine, Acetylcholine and NPY in aortas supernatant from *Ldlr^−/−^ll17rc^WT^* and *Ldlr^−/−^ Il17rc*^Δ^*^IEC^* (n=4-13) mice as determined by ELISA. ∗p<0.05. Student’s t-test.

To further extend our observation that upregulated neuronal signature was present in aortas along with inflammatory signature, we found neuronal expansion and a significant increase of NF200 (pan-neuronal), TH (adrenergic) and NPY (sympathetic) neurons in aortic roots of *Ldlr^−/−^ll17rc*^ΔIEC^ mice (Figure 1G, H). 3D Lightsheet microscopy imaging of whole mount atherosclerotic aortas cleared with iDISCO further confirmed expansion of TH^+^ sympathetic neuronal axons in *Ldlr^−/−^ Il17rc*^Δ*IEC*^ as compared to *Ldlr^−/−^ll17rc*^WT^ mice (Figure 1I). In line with heightened neuronal expansion, we observed elevated levels of adrenergic transmitters Norepinephrine and NPY, but not Acetylcholine in aortas of *Ldlr^−/−^ll17rc*^ΔIEC^ mice (Figure 1J).

Altogether, these data suggest previously underappreciated site- and cell-type specific role of intestinal IL-17RC signaling in atherosclerosis implying an important connection between IL-17RC signaling in the gut and neuronal and inflammatory pathways in atherosclerotic vessel, which culminates in augmented atherosclerosis.

### IL-17RC signaling in IEC controls barrier funtion and inflammation in the intestine

To understand how IL-17RC signaling in IEC may impact atherosclerosis development, we analyzed changes in the intestines of WD-fed *Ldlr^−/−^ll17rc*^Δ*IEC*^ and *Ldlr^−/−^ll17rc^WT^*mice. Histological analysis of ileum revealed reduction of mucus as determined by Muc2 staining and overall number of mucus-producing Goblet cells and junction barrier protein Claudin 1 (Figure 2A, B) in *Ldlr^−/−^ll17rc*^ΔIEC^ mice. PAS staining also demonstrate a reduction of mucus layer as well as numbers of Paneth cells (Figure S2A). Transcriptional analyses revealed upregulation of multiple inflammatory genes including *Il17a*, *Il1a, Il1b*, *Lcn2, Ifng* in ileum of WD-fed *Ldlr^−/−^ Il17rc*^ΔIEC^ mice compared to *Ldlr ^−/−^ll17rc*^WT^ controls; while genes encoding for antimicrobial peptides, hypoxia response and lipid metabolism (Figure 2C, D) as well as tight junction proteins (TJP) were downregulated (Figure S2B). Consequently, Ingenuity pathway analysis (IPA) revealed strong upregulation of IL-17, IFN, LPS and uric acid upstream regulators, while regulators for IL-10RA, butiric acid and AHR pathways were donwregulated (Figure S2C). In line with reduction of barrier proteins, *Ldlr^−/−^ll17rc*^ΔIEC^ mice showed increased intestinal permeability as determined by FITC-dextran translocation (Figure 2E).

**Figure 2.**
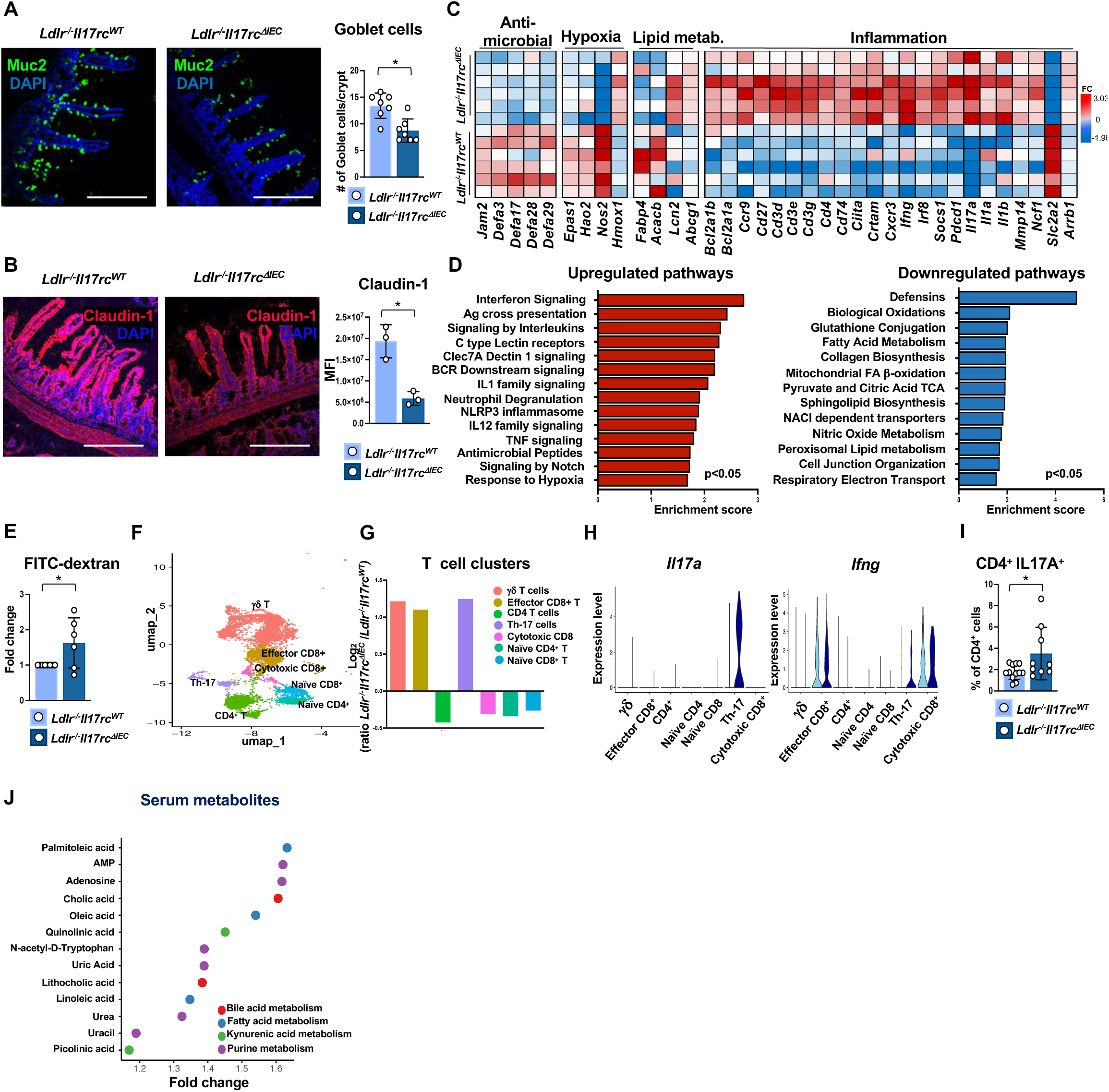
Intestinal IL-17RC deficiency reduces barrier function, enhances intestinal inflammation and alters microbiota. **A**. Representative immunofluorescent images showing Mucin (Mucin2; green) and DAPI (blue) staining of ileum sections (scale bar =100 μm) and quantification of Mucin2^+^ goblet cells numbers in *Ldlr^−/−^ll17rc^WT^* and *Ldlr^−/−^ll17rc*^Δ^*^IEC^* mice (n=7) after 16 weeks of WD-feeding. *p<0.05, Data are mean ± SEM from 3 independent experiments. Student’s t-test**. B**. Representative immunofluorescence images showing Claudin-1 (red) and DAPI (blue) staining of ileum sections (scale bar =100 μm) and quantification of Claudin-1 protein level of *Ldlr^−/−^ll17rc^WT^* and *Ldlr^−/−^ll17rc*^Δ^*^IEC^* mice (n=3). Data are mean ± SEM from 3 independent experiments. *p<0.05, Student’s t-test. RNA-seq analysis of differentially expressed genes in Ileums of *Ldlr^−/−^ll17rc^WT^* and *Ldlr^−/−^ll17rc*^Δ^*^IEC^* mice with a heatmap (**C**) (FC-Fold changes) and Ingenuity Pathway Analysis (IPA) (**D**). **E**. Fluorescent FITC-dextran measurements in the serum of *Ldlr^−/−^ll17rc^WT^* and *Ldlr^−/−^ll17rc*^Δ^*^IEC^* (n=6) mice as indication of intestinal permeability, *p<0.05, Student’s t-test. Single cell RNA sequencing analysis (10X Genomics Chromium) of CD45^+^ cells from Ileum tissue from WD-fed separately housed *Ldlr^−/−^ll17rc^WT^* and *Ldlr^−/−^ll17rc*^Δ^*^IEC^* mice. Cells were clustered by differential gene expression. **F.** UMAP plot showing clusters of T cells identified by gene expression profiles. **G.** Normalized cell numbers representing upregulated (positive Log2Foldchange) and downregulated (negative Log2Foldchange) clusters among T cells in ileum of *Ldlr^−/−^ll17rc*^Δ^*^IEC^* and *Ldlr^−/−^ll17rc^WT^* mice. **H.** Violin plots of selected differentially expressed genes among T cells in *Ldlr^−/−^ll17rc*^Δ^*^IEC^* vs *Ldlr^−/−^ll17rc^WT^* mice. **I.** FACS analysis of IL-17A expression in CD4^+^ T cells from ileum of *Ldlr^−/−^ll17rc^WT^* and *Ldlr^−/−^ll17rc*^Δ^*^IEC^* mice (n=9-11), ∗p<0.05. Student’s t-test. **J.** Dotplot of significantly changed serum metabolites in separately housed *Ldlr^−/−^ll17rc^WT^* and *Ldlr^−/−^ll17rc*^Δ^*^IEC^* mice fed with WD for 16 weeks. Representative data from 3 independent metabolomic analyses.

Since we detected barrier perturbation and increased translocation of microbial products, we next examined specific changes in intestinal immune cell composition and function. We conducted scRNA-seq of FACS sorted CD45^+^ cells from intraepithelial lymphocyte (IEL fraction) and lamina propria layer lymphocyte (LPL fraction) of ileums of WD-fed, separately housed *Ldlr^−/−^ll17rc*^Δ*IEC*^ and *Ldlr^−/−^ll17rc^WT^* mice and performed clustering to identify distinct cell populations in combined fractions (Figure 2F and Figure S2D, E, F, G). We found a significant increase in γδ T cells, Th17 CD4^+^ and effector CD8 T cells, while naive T cells were reduced (Figure 2G). Importantly, Th17 (CD4^+^) cells were the only producers of IL-17A as determined by scRNA-seq and confirmed by FACS (Figure 2H, I), while at least several cell subsets contributed to IFNγ production. Overall, our data showed that IEC-specific ablation of IL-17RC resulted in altered intestinal barrier function and increased inflammatory activation in the intestine of atherosclerosis-bearing mice.

### IL-17RC signaling in IEC controls microbiota composition in mice with atherosclerosis

Unhealthy diets, intestinal inflammation and altered barrier functions were previously associated with changes in intestinal microbiota. Next, to determine if enhanced atherosclerosis in *Ldlr^−/−^ Il17rc*^ΔIEC^ mice and heightened intestinal inflammation are associated with altered microbiota composition, we conducted shotgun metagenomic sequencing of DNA isolated from cecum luminal content. We found significant differences in overall luminal microbiome bacterial composition (Figure S3A) between separately housed, WD-fed *Ldlr^−/−^ Il17rc*^ΔIEC^ and *Ldlr^−/−^ll17rc^WT^*littermate mice.

As epithelium-bound and invasive bacteria may have disproportionally larger effect on immune activation or distant systemic physiological processes, we further analyzed the composition of adhesive bacteria in ileum, jejunum and colons from WD-fed *Ldlr^−/−^ Il17rc*^Δ*IEC*^ and *Ldlr^−/−^ll17rc^WT^* atherosclerosis-bearing mice by 16SRNA-seq of washed tissues. We found substantial differences in overall adhesive microbiome composition between IL-17RC deficient and sufficent mice with bacteria overrepresented in epithelium being different from those overrepresented in luminal compartment.

To determine if changes in microbiome composition in WD-fed *Ldlr^−/−^ Il17rc*^Δ*IEC*^ mice together with weakened barrier function may further lead to changes in circulating systemic metabolites, we conducted serum metabolomics. Metabolites from purine, fatty acids, bile acids and kynorenic acid metabolisms pathways were elevated in WD-fed *Ldlr^−/−^ Il17rc*^Δ*IEC*^ atherosclerosis-bearing mice, including significant upregulation of uric acid (purine metabolism), oleic, palmitoleic and linoleic acids (fatty acid metabolism), cholic and litocholic acids (bile acid metabolism) (Figure 2J). Althogether, disruption of IL-17RC in intestinal epithelium leads to significant changes in luminal and epithelium-adhesive microbiota and systemic increase in distinct metabolites.

### Enhanced aortic inflammation and expansion of peripheral neurons in atherosclerosis are functionally dependent on microbiota

We next sought to examine if altered microbiota composition is directly and mechanistically responsible for enhanced atherosclerosis in *Ldlr^−/−^ll17rc*^Δ*IEC*^ mice. We first co-housed *Ldlr^−/−^ Il17rc*^Δ*IEC*^ and *Ldlr^−/−^ll17rc^WT^* littermate mice to ensure natural way of microbiota transfer and equalization. In contrast to separately housed mice (Figure 1A and B), no significant differences in atherosclerotic lesion sizes were detected (Figure 3A). Furthermore, ablation of microbiota by broad-spectrum antibiotics reduced the disease in *Ldlr^−/−^ll17rc*^Δ*IEC*^ mice, abrogating the difference with controls; but did not significantly affect lesion sizes in *Ldlr^−/−^ll17rc^WT^*mice (Figure 3B and C). Interestingly, *Ldlr^−/−^ll17rc^WT^*mice developed bigger lesions when co-housed with *Ldlr^−/−^ Il17rc*^Δ*IEC*^ mice suggesting that transfer of potentially pathogenic microbiota from mice with IEC-specific IL-17RC ablation worsens the disease in *Ldlr^−/−^ll17rc*^WT^ controls (Figure 3C). These altogether implicates microbiota changes as a direct driver of accelerated atherosclerosis in *Ldlr^−/−^ Il17rc*^Δ*IEC*^ mice.

**Figure 3.**
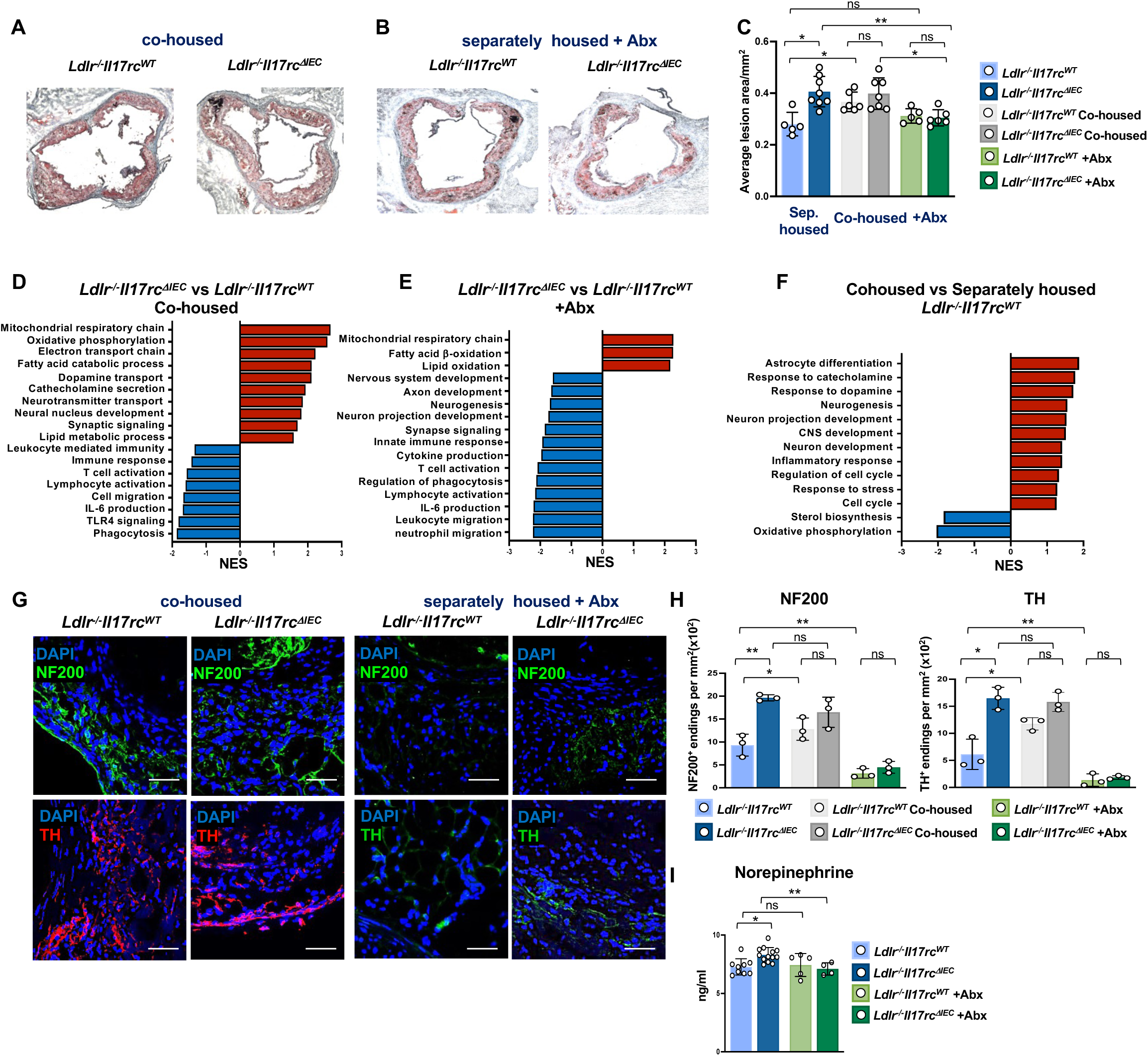
IL-17RC-mediated alterations of microbiota drive atherosclerosis and neuronal expansion in the aortas. **A.** Representative images of aortic root sections from WD-fed, co-housed littermate *Ldlr^−/−^ll17rc^WT^* and *Ldlr^−/−^ll17rc*^Δ^*^IEC^* mice and (**B**) separately housed mice with microbiota depleted by broad spectrum antibiotics (+ABX) after 16 weeks WD-feeding. **C**. Quantitative comparison of atherosclerotic lesion size of *Ldlr^−/−^ll17rc^WT^* and *Ldlr^−/−^ll17rc*^Δ^*^IEC^* mice in (**A**) and (**B**) (n = 5-8). Data are mean ± SEM from at least 3 independent experiments. *p<0.05, **p<0.01. Two-way ANOVA followed by Tukey’s post-hoc test was applied. Gene Set Enrichment Analysis (GSEA) of GO Biological Processes (GOBP) from bulk RNA-seq of aortas from co-housed *Ldlr^−/−^ll17rc^WT^* and *Ldlr^−/−^ Il17rc*^Δ^*^IEC^* mice (**D**), +ABX *Ldlr^−/−^ll17rc^WT^* and *Ldlr^−/−^ll17rc*^Δ^*^IEC^* mice (**E**) and co-housed and separately housed *Ldlr^−/−^ll17rc^WT^* mice (**F**). **G.** IF representative image of aortic root sections from *Ldlr^−/−^ll17rc^WT^* and *Ldlr^−/−^ll17rc*^Δ^*^IEC^* co-housed mice or +ABX mice stained for NF200 (top) and TH (bottom), scale bar is 50 μm. **H**. Quantitative comparison of NF200+ axons (n = 3) and TH+ axons (n = 3) in separately housed (from Figure 1), co-housed and +Abx *Ldlr^−/−^ll17rc^WT^* and *Ldlr^−/−^ll17rc*^Δ^*^IEC^* mice. Data are mean ± SEM from at least 3 independent experiments. *p<0.05, **p<0.01. Two-way ANOVA followed by Tukey’s post-hoc test was applied. **I**. Serum Norepinephrine level in separately housed (from Figure1) and +ABX *Ldlr^−/−^ll17rc^WT^* and *Ldlr^−/−^ll17rc*^Δ^*^IEC^* mice (n=4-13) determined by ELISA. Data are mean ± SEM from at least 3 independent experiments. *p<0.05, **p<0.01. Two-way ANOVA followed by Tukey’s post-hoc test was applied.

If enhanced atherosclerosis phenotype is transferrable by and dependent on microbiota, then co-housing would results in acquisition of “potentially pathogenic” species by *Ldlr^−/−^ll17rc^WT^* mice. Therefore, we next asked whether changes in microbiota composition regulate inflammation and neuronal outgrowth in aorta. We analyzed neuronal and inflammatory gene expression in co-housed *Ldlr^−/−^ll17rc^WT^* and *Ldlr^−/−^ll17rc*^Δ*IEC*^ mice and in microbiota-depleted mice in comparison with separately housed cohorts. We found that neuronal and inflammatory signatures were indeed microbiota dependent; and were upregulated in *Ldlr^−/−^ll17rc^WT^* mice co-housed with *Ldlr^−/−^ Il17rc*^Δ*IEC*^ mice (Figure 3D-F). Moreover, both inflammatory and neuronal signatures were significantly diminished in aortas of both *Ldlr^−/−^ll17rc^WT^* and *Ldlr^−/−^ll17rc*^Δ*IEC*^ mice when microbiota was depleted by antibiotics (Figure 3E). Immunofluorescent (IF) imaging of aortic roots revealed elevated presence of NF200^+^ neurons and TH^+^ adrenergic neurons in *Ldlr^−/−^ll17rc^WT^* mice co-housed with *Ldlr^−/−^ll17rc*^Δ*IEC*^, while microbiota depletion significantly reduced the amount of neuronal axons (Figure 3G and H) as well as levels of circulating Norepinephrine (Figure 3I). These data suggest causal connection between deregulated intestinal microbiota and enhanced neuronal growth and inflammatory activation in aorta.

### Loss of Intestinal IL-17RC facilitates IL-17^+^ γδ T cell accumulation in atherosclerotic aorta

Accumulation of various immune and inflammatory cells in the aortas accompanies atherosclerosis development and progression. scRNA-seq of aortic CD45^+^ cells (Figure S4A-E) revealed heightened accumulation of several types of T cells in aortas of *Ldlr^−/−^ll17rc*^Δ*IEC*^ with the most notable increase coming from γδ T cells (Figure 4A and B). Furthermore, differential gene expression (DEG) analysis showed that aortic γδ T cells represent a unique population expressing *II17a*, which was increased in *Ldlr^−/−^ll17rc*^Δ*IEC*^ mice (Figure 4C). Furthermore, aortic γδ T cells also expressed higher level of *Tnf*, *Itgae* ((CD103), involved to γδ T cell recruitment) as well as high levels of *Cxcr6*, while almost lacked *Ifng* expression (Figure 4C). We also found that γδ T cells expressed *Gpr65*, recently reported receptor for oleic acid^28^, a metabolite elevated in *Ldlr^−/−^ Il17rc*^Δ*IEC*^ mice (Figure 4C). We further confirmed increased accumulation of γδ T cells and IL-17A production in atherosclerotic aortas from separately housed *Ldlr^−/−^ll17rc*^Δ*IEC*^ mice by intracellular cytokine staining and FACS (Figure 4D, E). Analysis of scRNA-seq dataset (GSE252243)^29^ of human vessels from patients with atherosclerosis also demonstrated presence of γδ T cells that express *IL17A* and *GPR65* (Figure 4F). Immunohistochemistry staining of human atheroma confirmed presence of γδ T cells and high degree of sympathetic TH^+^ innervation in human atherosclerosis (Figure 4G).

**Figure 4.**
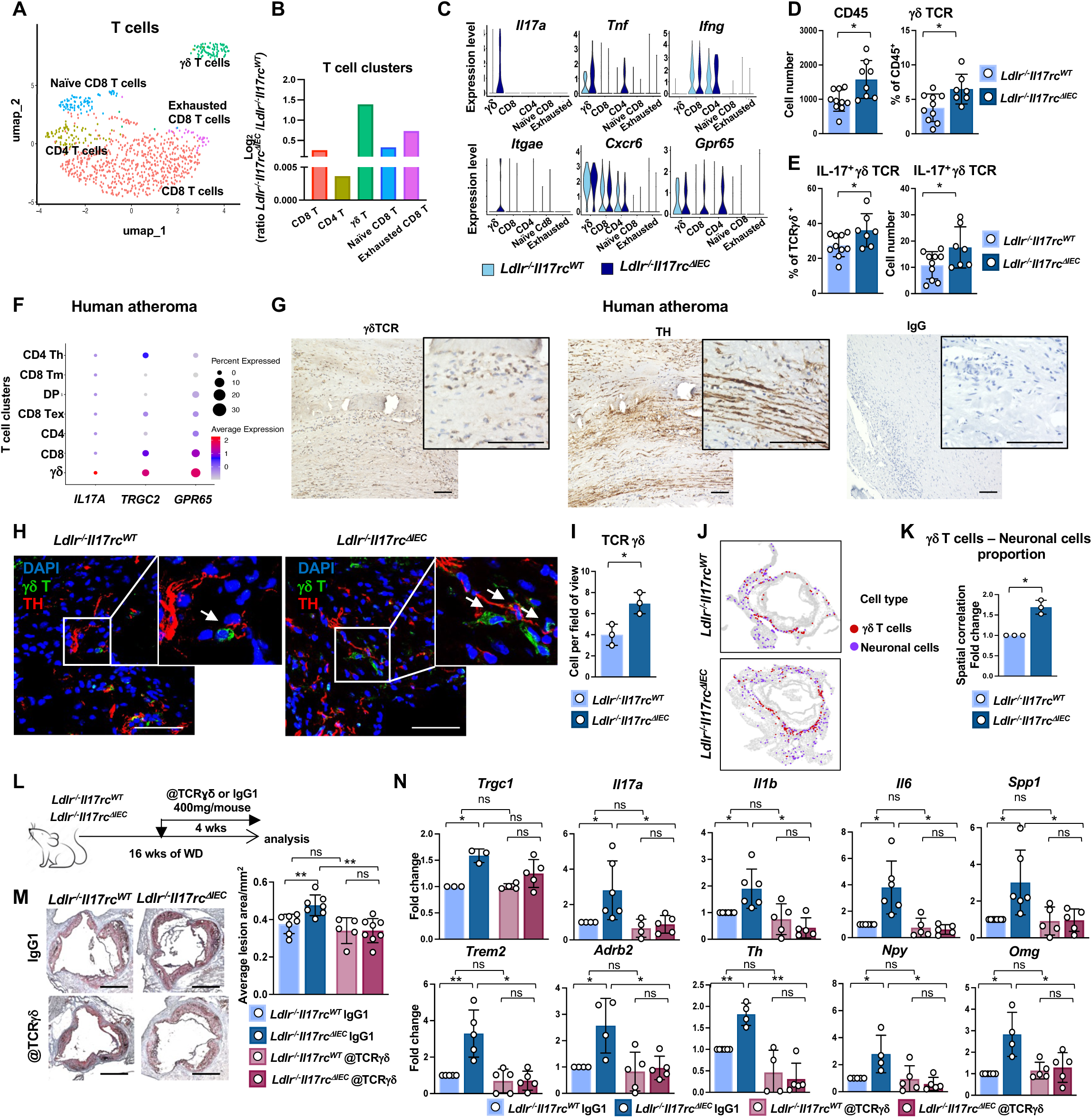
Intestinal IL-17RC deficiency results in accumulation of IL-17^+^ γδ T cells in atherosclerotic aorta that promotes neuronal expansion and atherosclerosis. Single cell RNA-seq analysis of FACS-sorted aortic CD45^+^ cells. **A.** UMAP of T cells clusters and normalized cell numbers among T cells (**B**) in aortas from *Ldlr^−/−^ll17rc^WT^* and *Ldlr^−/−^ll17rc*^Δ^*^IEC^* mice. **C.** Violin plots of selected differentially expressed genes between T cells from aortas of *Ldlr^−/−^ll17rc*^Δ^*^IEC^* and *Ldlr^−/−^ Il17rc^WT^* mice. FACS analysis of total CD45^+^ cells, γδ T cells (**D**) and IL-17 produced by γδ T cells (**E**) in aorta from *Ldlr^−/−^ll17rc^WT^* and *Ldlr^−/−^ll17rc*^Δ^*^IEC^* mice (n=7-10). Data are mean ± SEM from at least 3 independent experiments. *p<0.05. Student’s t-test. **F**. Single Cell RNA sequencing analysis of human asymptomatic atherosclerosis plaque (GSE252243 dataset). Annotations are derived from cluster specific analysis. Dotplot of T cells clusters from scRNA seq of human atherosclerotic plaque expressing *IL17A, TRGC2, GPR65*. **G**. Histological images of human atheroma stained for TCR γδ, TH and isotype control. Scale bar is 200 μm. **H** .IF images of aortic root sections from *Ldlr^−/−^ll17rc^WT^* and *Ldlr^−/−^ll17rc*^Δ^*^IEC^* mice stained TH and γδ TCR. Scale bar is 50 μm. **I**. Quantitative comparison of γδ T cells between *Ldlr^−/−^ll17rc^WT^* and *Ldlr^−/−^ll17rc*^Δ^*^IEC^* (n = 3) mice. Data are mean ± SEM from at least 3 independent experiments. *p<0.05, Student’s t-test. **J**. Representative Xenium spatial transcriptomic images of the aortic root from *Ldlr^−/−^ll17rc^WT^* and *Ldlr^−/−^ll17rc*^Δ^*^IEC^* mice demonstrate distinct co-localization of γδ T cells and neuronal cells. **K**. Quantitative comparison of γδ T cells – Neuronal cells proportion between *Ldlr^−/−^ll17rc^WT^* and *Ldlr^−/−^ll17rc*^Δ^*^IEC^* (n = 3) mice. *p<0.05, Student’s t-test. **L.** Scheme of the experiment. *Ldlr^−/−^ll17rc^WT^* and *Ldlr^−/−^ll17rc*^Δ^*^IEC^* mice were administered with anti-gd TCR antibody for 4 weeks starting at 12 weeks of WD feeding. **M**. Representative images of aortic root sections and quantitative comparison of atherosclerotic lesion sizes between *Ldlr^−/−^ Il17rc^WT^* and *Ldlr^−/−^ll17rc*^Δ^*^IEC^* mice treated with anti-γδ TCR or isotype control (n=5-8). **N.** Expression of selected inflammatory and neuronal genes in the aortas from *Ldlr^−/−^ll17rc^WT^* and *Ldlr^−/−^ll17rc*^Δ^*^IEC^* mice treated with anti- γδ TCR antibody or isotype control (n=4-6). Gene expression was normalized to *Rpl32* and then to gene expression in *Ldlr^−/−^ll17rc^WT^* treated with IgG1. Data are mean ± SEM from at least 3 independent experiments. *p<0.05, **p<0.01. Two-way ANOVA followed by Tukey’s post-hoc test was applied.

We next examined if T cells, and particularly γδ T cells, are co-localized with neurons. We co-stained nerves with γδTCR T cell markers in aortic root sections of separately housed *Ldlr^−/−^ Il17rc*^Δ*IEC*^ and *Ldlr^−/−^ll17rc^WT^* mice fed with WD for 16 weeks. Consistent with our scRNA-seq and FACS results, we found more γδ T cells in aortas of *Ldlr^−/−^ll17rc*^Δ*IEC*^ mice, and furthermore observed that some γδTCR^+^ cells are located in a close proximity to TH^+^ sympathetic neurons (Figure 4H and I) suggesting possible link between IL-17A^+^ γδ T cells and neurons in the vessel wall. Xenium spatial transcriptomics analysis revealed increased co-localization between γδ T cells and neurons suggesting their interactions in aortas of *Ldlr^−/−^ll17rc*^Δ*IEC*^ mice (Figure 4J and K). To examine the mechanistic link between elevated presence of γδ T cells and expanded neuronal axons *in vivo*, we administered non-depleting, blocking anti-TCRγδ antibody (UC7-13D5) into separately housed *Ldlr^−/−^ll17rc*^Δ*IEC*^ and *Ldlr^−/−^ll17rc^WT^*mice for the last 4 weeks of WD feeding (Figure 4L). We found that functional γδ T cell blockade significantly reduced atherosclerotic lesions in *Ldlr^−/−^ll17rc*^Δ*IEC*^ but not *Ldlr^−/−^ll17rc^WT^*mice as compared to IgG treated controls attributing the differences in the disease to γδ T cell accumulation (Figure 4M). Transcriptional analysis of isolated aortas revealed that γδ TCR blockade did not affect *Trgc1* expression, and therefore, presence of γδ T cells (Figure 4N), but significantly reduced inflammatory (*Il17a, Il1b, Il6, Spp1, Trem2)* and neuronal *(Th1, Npy, Omg)* gene expression (Figure 4N). These data suggest that aortic IL-17A^+^ γδ T cell accumulation directly modulates neuronal outgrowth and activation as well as inflammation, contributing to heightened atherosclerosis in *Ldlr^−/−^ll17rc*^Δ*IEC*^ mice.

### Augmented atherosclerosis depends on cytokine-mediated regulation of neurons and their activity

To further establish whether augmented atherosclerosis development in WD-fed *Ldlr^−/−^ll17rc*^Δ*IEC*^ mice is directly driven by neuronal expansion, we performed chemical denervation of sympathetic neurons by administering *Ldlr^−/−^ll17rc*^Δ*IEC*^ and *Ldlr^−/−^ll17rc^WT^* mice with 6-Hydroxydopamine (6-OHDA). We found that 6-OHDA substantially reduced the disease in both groups of mice (Figure 5A, B) and the effect was more pronounced in *Ldlr^−/−^ll17rc*^Δ*IEC*^ mice and, 6-OHDA abolished difference between *Ldlr^−/−^ll17rc*^Δ*IEC*^ mice and controls. This was further accompanied by downregulation of inflammatory (*Il6, Spp1, Trem2*) and neuronal (*Th1, Dbh, Npy*) gene expression in the aortas of *Ldlr^−/−^ll17rc*^Δ*IEC*^ mice (Figure 5C), indicating that these pathways in atherosclerosis are at least in part controlled by sympathetic neurons.

**Figure 5.**
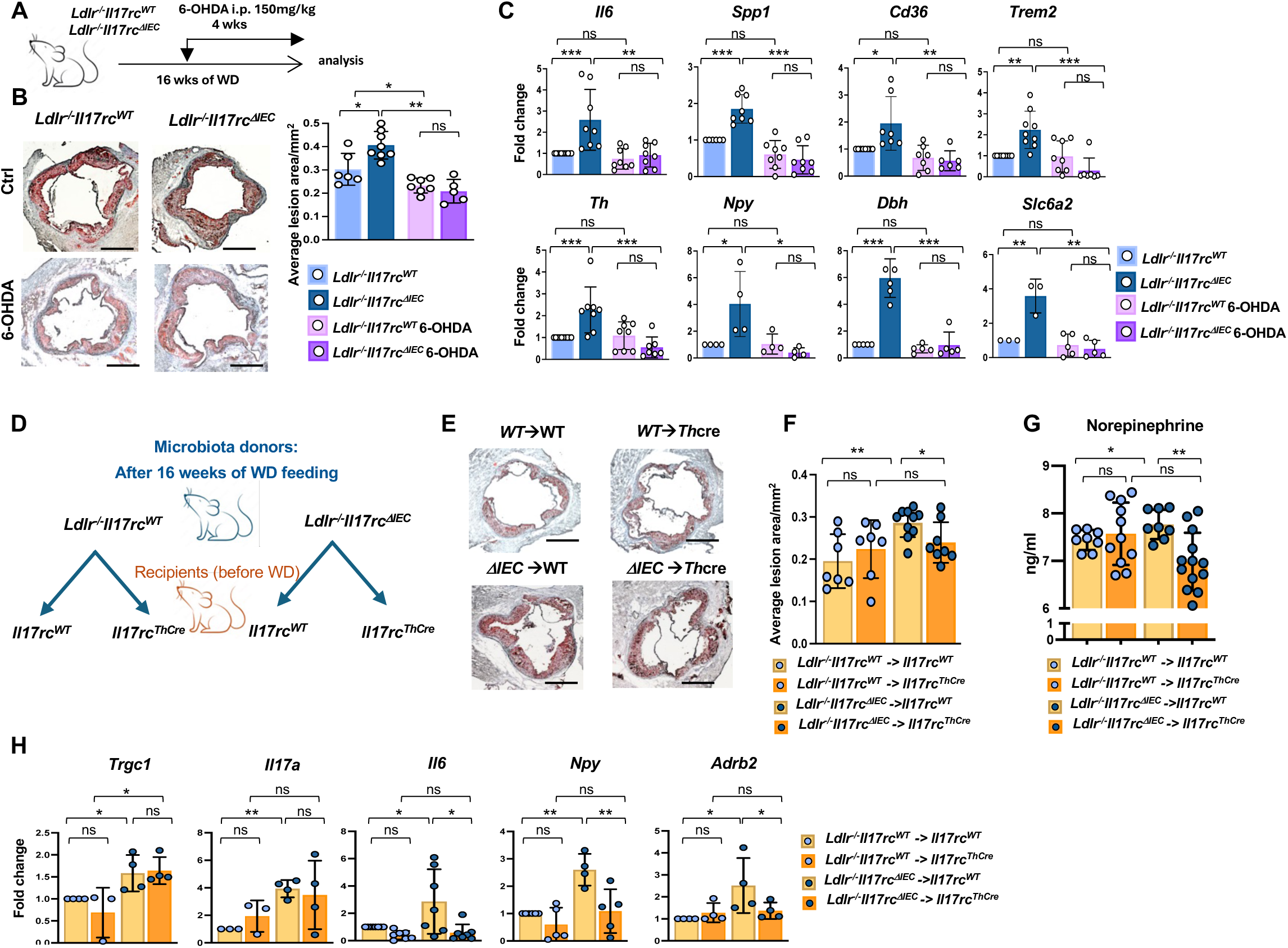
IL-17RC signaling in neurons mediates microbiota-driven neuro-inflammatory responses in aorta and promotes atherosclerosis. **A**. Scheme of the experiment. *Ldlr^−/−^ll17rc^WT^* and *Ldlr^−/−^ll17rc*^Δ^*^IEC^* mice were administered with 6-OHDA for 4 weeks starting at 12 weeks of WD feeding. **B**. Representative images of aortic root sections and quantitative comparison of atherosclerotic lesion sizes between *Ldlr^−/−^ll17rc^WT^* and *Ldlr^−/−^ll17rc*^Δ^*^IEC^* mice treated with 6-OHDA or control (n=5-8). Data are mean ± SEM from 3 independent experiments. *p<0.05, **p<0.01. Two-way ANOVA followed by Tukey’s post-hoc test was applied. **C**. Expression of selected Inflammatory and neuronal genes in the aortas from *Ldlr^−/−^ll17rc^WT^* and *Ldlr^−/−^ll17rc*^Δ^*^IEC^* mice treated with 6-OHDA or control (n=4-8). Gene expression was normalized to *Rpl32* and then to gene expression in *Ldlr^−/−^ll17rc^WT^*. Data are mean ± SEM from 3 independent experiments. *p<0.05, **p<0.01, two-way ANOVA followed by Tukey’s post-hoc test was applied. **D.** Scheme of microbiota transfer experiment. **E.** Representative images of aortic root sections and (**F**) quantitative comparison of atherosclerotic lesion size between mice after microbiome transfer in *Ldlr^−/−^ll17rc^WT^* ➔ *ll17rc^WT^*, *Ldlr^−/−^ll17rc^WT^* ➔ *ll17r^ThCre^, Ldlr^−/−^ll17rc*^Δ^*^IEC^* ➔ *ll17rc^WT^* or *Ldlr^−/−^ll17rc*^Δ^*^IEC^* ➔ *ll17rc^ThCre^* (n = 7-10). Data are mean ± SEM from 3 independent experiments. *p<0.05, **p<0.01, two-way ANOVA followed by Tukey’s post-hoc test was applied. **G.** Serum Norepinephrine from FMT *Ldlr^−/−^ll17rc^WT^* ➔ *ll17rc^WT^*, *Ldlr^−/−^ll17rc^WT^* ➔ *ll17r^ThCre^, Ldlr^−/−^ll17rc*^Δ^*^IEC^* ➔ *ll17rc^WT^* or *Ldlr^−/−^ll17rc*^Δ^*^IEC^* ➔ *ll17rc^ThCre^* mice (n=4-6) as determined by ELISA. Data from 3 independent experiments, *p<0.05, **p<0.01, two-way ANOVA followed by Tukey’s post-hoc test was applied. **H.** Expression of selected Inflammatory and neuronal genes in the aortas from *Ldlr^−/−^ll17rc^WT^ Il17rc^WT^*, *Ldlr^−/−^ll17rc^WT^* ➔ *ll17r^ThCre^, Ldlr^−/−^ll17rc*^Δ^*^IEC^* ➔ *ll17rc^WT^* or *Ldlr^−/−^ll17rc*^Δ^*^IEC^* ➔ *ll17rc^ThCre^* mice (n=4-7) mice. Gene expression was normalized to *Rpl32*. Data are mean ± SEM from 3 independent experiments, ∗p<0.05, **p<0.01, two-way ANOVA followed by Tukey’s post-hoc test was applied.

Neurons express cytokine receptors; and thus their growth, survival or function can be regulated by cytokines^19^. Next, we isolated neurons from multiple ganglia along spinal cord, including nodose root ganglia known to contain bodies of neurons innervating aortas, and stimulated them *ex vivo* with recIL-17A. Transcriptional analysis revealed that IL-17A enhanced expression of genes associated with inflammation (*Spp1, Csf2, Cxcl5, Ifi208, Cd40*), neuronal interaction (*Hey2, Lif, Ccl25*), tissue repair and canonical type 17 pathway (Figure S5A); indicating that direct neuronal IL-17A signaling may cause transcriptional shifts leading to increased innervation, induction of inflammatory signaling and peripheral neuron-dependent augmentation of atherosclerosis. Next, we tested whether neuron-specific IL-17RC signaling is required for the expansion and activation of neurons and atherosclerosis in WD-fed *Ldlr^−/−^ll17rc*^Δ*IEC*^ with dysbiosis and heightened gut inflammation. We generated *Il17rc* THCre mice that lack IL17RC in sympathetic neurons^30^, however, these mice could not be further crossed onto *Ldlr^−/−^* background because these genes are located on the same chromosome. We therefore administered *Il17rc* THCre mice with LDLR-CRISPR AAV to ablate LDLR expression in the liver and induce atherosclerosis, followed by 16 weeks of WD feeding ^31^. No difference in atherosclerosis development was observed in *Il17rc* THCre^−^ versus *Il17rc* THCre^+^ mice (Figure S5B, C), indicating that this pathway is not essential in the absence of intestinal barrier and microbiota perturbations. Since pro-atherogenic phenotype was transferrable by microbiota (Figure 3), we performed fecal transplant from atherosclerosis-bearing separately housed WD-fed *Ldlr^−/−^ Il17rc*^Δ*IEC*^ and WD-fed *Ldlr^−/−^ll17rc^WT^* mice to *ll17rc* THCre^−^ and *Il17rc* THCre^+^ recipients, administered them with LDLR-CRISPR-AAV to induce atherosclerosis, and fed with WD for 16 weeks (Figure 5D). Analysis of atherosclerotic lesions revealed that microbiota transfer from *Ldlr^−/−^ll17rc^WT^* mice did not alter atherosclerosis development, while microbiota from *Ldlr^−/−^ll17rc*^Δ*IEC*^ mice significantly induced the disease in *Il17rc* THCre^−^ mice (Figure 5E, F). However, ability of diseased microbiota to promote atherosclerosis was significantly blunted in *Il17rc* THCre^+^ mice lacking IL-17RC signaling in sympathetic neurons (Figure 5E, F). Serum norepinephrine was significantly elevated in FMT *Ldlr^−/−^ll17rc*^Δ*IEC*^ *->Il17rc* THCre^−^ mice as compared to *Ldlr^−/−^ Il17rc^WT^ ->Il17rc* THCre^−^ controls, but strongly reduced in FMT *Ldlr^−/−^ll17rc*^Δ*IEC*^ *->Il17rc* TH Cre^+^ mice which correlated with reduced atherosclerotic lesion sizes (Figure 5G). Transcriptional analysis of isolated aortas showed that FMT from *Ldlr^−/−^ll17rc*^Δ*IEC*^ simulated expression of *Trgc1, Il17a, Il6*, adrenergic receptor *Adrb2* as well as sympathetic neuronal marker *Npy* in *ll17rc* TH Cre^−^ recipients. While *Trgc1* and *Il17a* expression was similar in *Ldlr^−/−^ll17rc*^Δ*IEC*^ *->Il17rc* TH Cre^+^ mice, we observed strong downregulation of *Il6, Ifnb1, Npy* and *Adrb2* in FMT *Ldlr^−/−^ll17rc*^Δ*IEC*^ *-> Il17rc* TH Cre^+^ aortas (Figure 5H). These data suggest that IL-17RC ablation in sympathetic neurons reduced their activation and suppressed inflammatory gene expression in the aortas, thereby resulting in at least partial protection from atherosclerosis (Figure 5E, F). These data altogether demonstrate that in the presence of intestinal barrier defects and/or dysbiotic microbiota, IL-17A is further induced distantly in aortas and its sympathetic neuron specific signaling is required for full scale neuronal expansion, induction of inflammation and progression of atherosclerosis.

### Sympathetic neurons influence pro-inflammatory macrophage activation in the aortas

Progressive atherosclerotic plaque growth instrumentally involves macrophages which take up lipids and cholesterol and form foam cells. To gain insights into the specific changes in aortic immune cell composition we further analyzed scRNA-seq dataset of FACS sorted, live CD45^+^ cells from aortas with atherosclerosis of separately housed *Ldlr^−/−^ll17rc*^Δ*IEC*^ and *Ldlr^−/−^ll17rc^WT^* mice fed with WD for 16 weeks (Figure S4A, B, C) which showed altered myeloid cells composition between *Ldlr^−/−^ll17rc^WT^* and *Ldlr^−/−^ll17rc*^Δ*IEC*^ mice (Figure S6A, B). Normalized cell ratio revealed an increase in representation of inflammatory IFNIC, Trem2^hi^-Slamf9 and Trem2^hi^-Gpnmb macrophages as well as classical monocytes in WD-fed *Ldlr^−/−^ll17rc*^Δ*IEC*^ mice, while TLR-CD209^low^, TLR-CD209^hi^ and MAC-AIR macrophages populations were reduced (Figure 6A, B). Differential gene expression analysis of IFNIC (interferon-driven) and Trem2^hi^-Slamf9 (“foamy macrophages”) clusters showed upregulation of pro-inflammatory pathways such as type I IFN response (in IFNIC) and IL-1β and IL-6 production (in Trem2^hi^-Slamf9) in *Ldlr^−/−^ll17rc*^Δ*IEC*^ mice (Figure 6C,D). FACS analysis of aortic immune cells confirmed an increase in overall macrophage numbers and particularly inflammatory IL-6-producing macrophages in WD-fed *Ldlr^−/−^ll17rc*^Δ*IEC*^ mice (Figure 6E). This altogether suggest that intestinal IL-17RC signaling regulates the emergence of pro-atherogenic macrophages in aorta.

**Figure 6.**
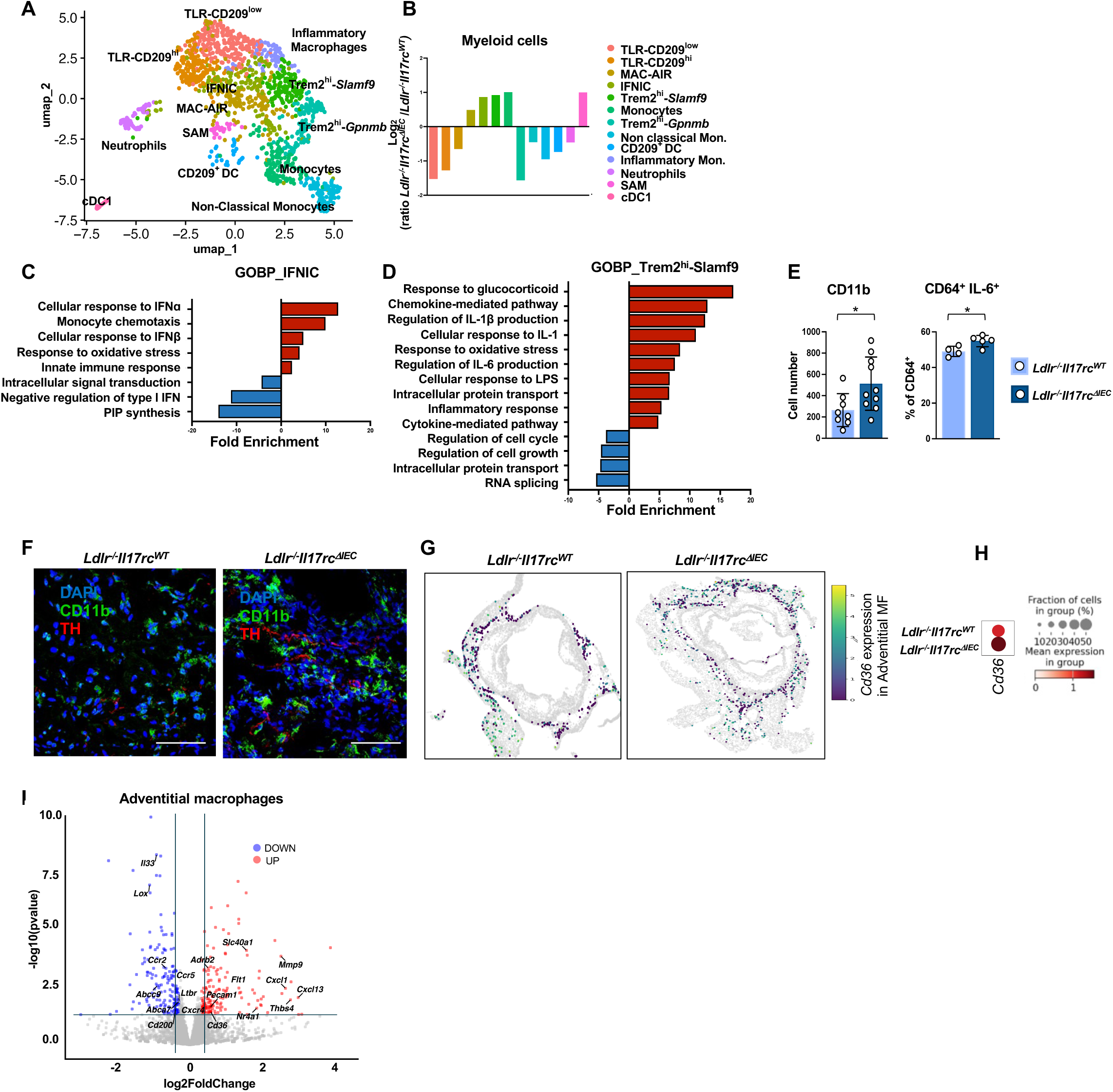
IL-17RC signaling deficiency in IEC participates in accumulation of inflammatory and foamy macrophages formation in atherosclerotic plaque. **A**. UMAP of myeloid cells clusters in aorta from *Ldlr^−/−^ll17rc^WT^* and *Ldlr^−/−^ll17rc*^Δ^*^IEC^* mice. **B.** Normalized cell numbers representing upregulated and downregulated clusters among myeloid cells in aortas from *Ldlr^−/−^ll17rc*^Δ^*^IEC^* and *Ldlr^−/−^ Il17rc^WT^* mice. Gene Ontology (GO) biological pathway (BP) enrichment analysis of differentially expressed genes in IFNIC **(C)** and Trem2^hi^-Slamf9 **(D)** macrophage clusters from *Ldlr^−/−^ll17rc^WT^* and *Ldlr^−/−^ll17rc*^Δ^*^IEC^* mice. **E.** FACS analysis of CD45^+^CD11b^+^ cells and IL-6 expression in CD64^+^CD11b^+^ cells from aortas of *Ldlr^−/−^ll17rc^WT^* and *Ldlr^−/−^ll17rc*^Δ^*^IEC^* mice (n=4-10). **F.** IF representative images of aortic root sections from *Ldlr^−/−^ll17rc^WT^* and *Ldlr^−/−^ll17rc*^Δ^*^IEC^* mice stained for CD11b (green), sympathetic neurons TH (red), DAPI (blue), scale bar=50μm. **G.** Representative spatial transcriptomic images of the aortic root from *Ldlr^−/−^ll17rc^WT^* and *Ldlr^−/−^ll17rc*^Δ^*^IEC^* mice show *Cd36* gene expression localized to adventitial macrophages. **H.** Quantification of *Cd36* gene expression in adventitial macrophages between *Ldlr^−/−^ll17rc^WT^* and *Ldlr^−/−^ll17rc*^Δ^*^IEC^* mice. **I.** Volcano plot of differentially expressed genes in adventitial macrophages between *Ldlr^−/−^ll17rc*^Δ^*^IEC^* and *Ldlr^−/−^ll17rc^WT^* mice. Only p<0.05 are shown.

Next, we tested whether sympathetic neurons could influence atherogenic macrophages. We observed expanded co-localization of macrophages next to Adrenergic neurons (TH) in aortic roots of mice with *Ldlr^−/−^ll17rc*^Δ*IEC*^ compared to control (Figure 6F). Spatial transcriptomics analysis revealed upregulation of *Cd36* in aortic macrophages, specifically those located in adventitia in *Ldlr^−/−^ll17rc*^Δ*IEC*^ mice compared to *Ldlr^−/−^ll17rc^WT^*control (Figure 6G, H). Furthermore, differential gene expression showed upregulation of *Adrb2* as well as pro-atherogenic *Cd36, Mmp9, Cxcl1, Pecam* genes in adventitial macrophages from *Ldlr^−/−^ll17rc*^Δ*IEC*^ mice, while anti-inflammatory *(Il33, Cd200)* and cholesterol reverse transport genes *(Abca7, Abcc9)* were downregulated (Figure 6I). *Trem2^+^* macrophages preferentially located in atherosclerotic plaque were not significantly affected by intestinal *Il17rc* deficiency (Figure S7A, B). Overall, our data suggest that activated sympathetic neurons via production of norepinephrine may stimulate inflammatory macrophage activation and lipid uptake, thereby contributing to atherosclerosis development.

Taken together our data demonstrate previously unappreciated cell-type specific roles of IL-17 in atherosclerosis and established that intestinal dysbiosis controlled by anti-atherogenic IL-17RC signaling in IEC drives the accumulation of IL-17 producing γδ T cells in the aortas. In contrast, in atherosclerotic aortas IL-17A stimulates activation and expansion of sympathetic neurons to promote neurogenesis, neuro-immune interactions, inflammatory macrophage activation and lipid uptake, thereby augmenting the disease.

## DISCUSSION

Atherosclerosis is a multifactorial disease driven by multiple risk factors including hyperlipidemia, genetics, environmental insults, obesity, diet, stress, smoking, sleep deprivation, type II diabetes and intestinal diseases such as IBD. Many of these factors are considered as “stand-alone contributors”, not necessarily as a part of bigger unifying mechanism. Only few of them – LDL/cholesterol management or inhibition of IL-1β pathway (CANTOS trial) along with dietary interventions are currently mainstream in clinical practice with beneficial effects, while new players like GLP1 agonists have not yet been tested.

Here we uncovered a multicellular systemic mechanism regulating atherosclerosis development which starts with suboptimal diet (WD) and deterioration of protective intestinal barrier and leads to a specific immune activation in aorta (γδ T cells; IL-17A) that promotes expansion and aberrant activation of sympathetic neurons. Enhanced aortic inflammation and neuronal expansion are strictly dependent on microbiota and this phenotype can be transferred by co-housing or reduced by microbiota depletion. Importantly neuronal presence and activity is required for further downstream regulation of aortic inflammation, macrophage activation, lipid uptake and atherosclerosis progression. This pathogenic sequence of events has multiple “entry points” to which various stimuli and atherosclerosis-risk factors can further feed into acceleration of the disease. For example, antibiotic abuse, dysbiosis and suboptimal diet can further fuel the cascade at the level of intestinal barrier. Stress via enhanced neuronal signaling would be able to induce even more inflammation and lipid uptake in aorta, provided that barrier-microbiota-neuronal circuits are already established. Finally, genetic or dietary-driven hyperlipidemia would result in pro-inflammatory macrophage activation, lipid uptake and plaque growth on top of inflammatory and neuronal signals already controlling this process.

We found that weakening of intestinal barrier (here specifically by virtue of cytokine signaling loss) leads to increased subclinical intestinal inflammation, changes in luminal and adhesive microbiota and circulating metabolites. Cytokine signaling in intestinal epithelial cells has been suggested to modulate microbiota composition and function^32^. While prior studies established an important role of IL-17A in regulation of mucosal immunity^18,33,34^; and IL-17RA ablation in the gut was linked to autoimmune diseases such as EAE due to enhanced IL-17A production ^18,35,36^, the role of IL-17RC signaling in intestinal epithelial cells as regulator of intestinal homeostasis and host-microbiota interactions in diet-driven atherosclerosis has never been investigated. Moreover, we previously found that elevated serum LPS and TMAO metabolite link altered gut permeability to macrophage activation and atherosclerosis^37,38^. Here we showed that IL-17RC signaling in intestinal epithelial cells plays a protective role in atherosclerosis due to its ability to control intestinal barrier and microbiota and, unexpectedly, sympathetic innervation in the aorta. The expansion of peripheral neurons in the aortas in humans and mice during atherosclerosis development had been recently demonstrated^39^. Our data suggest that expansion and activation of neurons in aortas is microbiota dependent. Fecal transfer from *Ldlr^−/−^ll17rc*^Δ*IEC*^ mice with developed atherosclerosis induced *Il17 and Il6* as well as sympathetic neuronal genes such as *Th1* and *Npy* in aortas suggesting athero-promoting roles for these bacteria.

How do these pro-atherogenic signals from intestine transmit to aortas? We found heightened inflammatory signature, and in particular, increased presence of IL-17A^+^ γδ T cells in the aortas. γδ T cells were detected in healthy vessels of mice and humans but their role in atherosclerosis is not fully understood. Our data demonstrate that γδ TCR blockade suppressed atherosclerosis in *Ldlr^−/−^ll17rc*^Δ*IEC*^ mice with altered microbiota but did not significantly affect the disease in *Ldlr^−/−^ Il17rc^WT^*controls, in line with earlier reports^40,41^. Activation of γδ T cells may be conferred by myeloid cells activated by bacterial ligands to produce IL-1β and IL-23 that in turn may regulate IL-17A production by γδ T cells^7,42–44^. It is also possible that aortic γδ T cells initially become activated in the intestine and are able to migrate to aorta. Presence of distinct bacteria was linked to host lipid metabolism changes and altered lipid and fatty acid absorption. Microbiota-dependent fatty acids, for examples short chain fatty acids (SCFA) were shown to suppress IL-17A production by intestinal γδ T cells^45^. Our studies revealed a significant increase in circulating monounsaturated fatty acids including oleic acid in serum of separately housed *Ldlr^−/−^ll17rc*^Δ*IEC*^ mice with heightened atherosclerosis. Furthermore, our data showed that aortic T cells, including γδ T cells express *Gpr65*, a recently described receptor for oleic acid^28^; and stimulation of γδ T cells with oleic acid induced *Il17a*, but not *Ifng* expression. Elevated circulating oleic acid had been previously correlated with presence of *Lactobacillus spp*^46^ and was associated with a greater risk of CVD^47^, similar to our observations. While role of oleic acid in human cardiovascular diseases is not unequivocal. Oleic acid was shown to promote vascular smooth muscle cells (VSMC) activation and re-programming of T cells toward Th9 cell differentiation *in vitro*^48,49^. Analysis of published human atheroma sequencing data revealed presence of γδ T cells in atherosclerotic plaque, that were characterized by *GPR65* and *IL17A* expression. Furthermore, histological staining revealed presence of γδ T cells and TH^+^ sympathetic neurons in human atheroma confirming our observations in a mouse model.

Notably, we found γδ T cells in a proximity to neurons suggesting possible connection between these cell types. Indeed, our data showed that γδ TCR pharmacological blockade reduced expression of sympathetic neuronal markers as well as inflammatory genes in the aortas. These data are in line with prior report that *Tcrd^−/−^* mice are characterized by reduced number of sympathetic neurons in the fat due to altered TGFβ production from parenchymal cells^50^. Overall, these observations suggest that microbiota and microbiota-dependent oleic acid may activate γδ T cells to produce IL-17A that could act locally in the vessel wall and stimulate local peripheral innervation.

Microbiota alteration was linked to neuroinflammation and altered social behavior^51^. However, while dysbiosis was linked to cardiovascular disease development, no studies to date establish the link between altered microbiota and enhanced sympathetic innervation in the vessel as potential mechanism of atherosclerosis. Microbiota-dependent IL-17A production was linked to autoimmune encephalomyelitis (EAE) (mouse model of multiple sclerosis)^18,52^. Furthermore, the direct effect of IL-17 signaling on neurons in the brain or skin had been previously suggested^22–24^. However, whether IL-17RC signaling may modulate activation of peripheral neurons in the aortas has not been investigated. We showed that genetic ablation of IL-17RC in sympathetic neurons was able to suppress microbiota-driven atherosclerosis in mice that received disease-promoting microbiota from atherosclerotic *Ldlr^−/−^ll17rc*^Δ*IEC*^ mice. This suggest that local upregulation of IL-17A in the diseased vessel maybe implicated to neuronal activation that in turn provides positive feedback loop regulating inflammation in atherosclerosis.

Macrophages represent a key immune cell population in atherosclerosis. The progression of the disease is always accompanied by accumulation of pro-inflammatory subsets and lipid-loaded foam cell formation^53^. We found an expansion of pro-inflammatory macrophage subsets in aortas of *Ldlr^−/−^ll17rc*^Δ*IEC*^ mice; and established that macrophages in the aortas co-localized with sympathetic neurons. We also confirm that aortic macrophages express Norepinephrine receptors in agreement with previous reports, and thus their function can be modulated by upregulation of sympathetic innervation resulted in Norepinephrine release^54^. While some types of innervation (vagus-to-spleen) results in decreased inflammation and less activated myeloid cells^55^, we observe that in aorta activity of sympathetic neurons promotes inflammation and pro-inflammatory macrophage accumulation that express more scavenger receptor CD36.

Studies in mice and humans showed that circulating levels of IL-17A correlate with atherosclerosis development, and accumulation of IL-17A producing T cells in atherosclerotic plaques was noted^8,9^. GWAS studies identified that polymorphisms in *IL17A* and its receptor *IL17RC* genes are associated with carotid artery thickness^56^. However, the role of IL-17 signaling in atherosclerosis remains controversial and mouse models utilizing global knockouts or systemic gain- or loss-of function approaches showed controversial effects^8–15^. The same knockout of IL-17A showed opposite results in different labs: while the acceleration of disease was found in *Il17a^−/−^Apoe^−/−^* mice^57^, a strong reduction in atherosclerosis in independently bred *Il17a^−/−^* or *Il17ra^−/−^Apoe^−/−^* mice was also reported^13^. Notably, most of earlier experiments were reported without specific mentioning of genetic background purity, housing conditions (in-house or acquired mice) or littermate control status complicating the interpretation of obtained results that did not account for IL-17 detrimental effects on microbiota. Our work uncovers that cell type specific IL-17 signaling may play opposing roles in atherosclerosis, solely dependent which cells respond to this cytokine. Pro-inflammatory properties of IL-17A in several inflammatory diseases and cancer made the field to swarm for studies trying to inhibit IL-17A, which successfully worked in psoriasis and arthritis^58^. However, while IL-17 was upregulated in patients with IBD^59^, its neutralization worsened the disease leading to the halt of clinical trials^60^, and IL-17 neutralization in psoriasis patients did not improve their CVD health despite psoriasis itself predisposes to CVD^61^. Moreover, IBD patients with enhanced inflammation and disrupted intestinal barrier have elevated risk of CVD and atherosclerosis despite being overall leaner than healthy control groups^62,63^. We found that epithelial IL-17RC signaling protects mice from atherosclerosis by regulating microbiota homeostasis and metabolite influx; while neuronal IL-17RC signaling in aorta is pro-atherogenic and stimulates neuronal outgrowth and activity, culminating in enhanced pro-inflammatory macrophage activation.

Taken together our data identified novel cell type specific role of IL-17 in atherosclerosis and demonstrate novel gut-peripheral nerves connection that control vessel wall neuro-immune interactions, inflammation and atherosclerosis.

## RESOURCE AVAILABILITY

### Lead contact

Request for further information and resources should be directed to and will be fulfilled by the Lead Contact, Ekaterina Koltsova

## ACKNOWLEDGMENTS

We acknowledge the help of Cedars-Sinai Medical Center facilities. We thank Dr. Daniel Mucida for critical reading of the manuscript and Dr. Ilya Kadnikov for constructive discussions. This work was partially supported by Cedars-Sinai Cancer funds to E.K.K. and S.I.G.; as well as NIH R01 HL173975 and R01 CA273925 grants to E.K.K.; R01 CA 287786 to S.I.G; furthermore R.R., A.K.D., and G.T. were supported in part by the Intramural Research Program of the NIH. T.Q.d.A.V. was supported in part by the NIH R01HL163908, NIH R01HL174008, R01DK128952, and R01DK138340 and K.E.J. was supported by Iris Cantor Women’s Health Center UCLA Clinical and Translational Science Institute pilot grant (UL1TR001881) and a UCLA SCORE on Cardiometabolic Health and Disease pilot grant (U54HL170326).

## AUTHORS CONTRIBUTION

A.M.M., S.I.G and E.K.K. designed the study, planned the experiments, analyzed the data. A.M.M., J.Z., M.T., K.N., P.N. and K.E.J. performed the experiments. A.M.M. and J.Z. performed scRNA experiments. A.M.M. and P.V. performed 10x Genomics Xenium in situ. J.A., S.K. and J.Z performed analysis of spatial transcriptomics. A.M.M. and M.C. performed iDISCO experiment. R.R. and A.K.D. performed microbiome analysis. B.S. provided advice on cytokine-producing cell recruitment. A.M.M. and A.K. performed bulk RNAseq and scRNA seq analyses. A.M.M., S.I.G., and E.K.K. wrote the manuscript with the input from other co-authors. S.I.G., E.K.K. and T.Q.d.A.V obtained funding and S.I.G. and E.K.K. supervised the work.

## DECLARATION OF INTERESTS

Authors declare no conflicts of interest

### Contact for reagents and resource sharing

Further information and requests for resources and reagents should be directed to and will be fulfilled by the Lead Contact, Ekaterina Koltsova

### Animals

All animal experiments were approved by the Animal Care Committee at Cedars Sinai Medical Center (CSMC). C57BL/6J control (000664), *Ldlr^−/−^* (002207), *Villin^Cre^* (021504), *Th^Cre^* (008601) were purchased from Jackson Laboratories and kept in our facility. *ll17rc^flox/flox^* (*ll17rc^flox/flox^* were generated in the lab in collaboration with Dr. Grivennikov (Figure S1 and ^27^). *ll17rc^flox/flox^* were bred with *Ldlr^−/−^*, *Villin^Cre^* or *Th^cre^* mice to obtain *Ldlr^−/−^* x *ll17rc^flox/flox^* x *Villin^Cre^* (referred in text as *Ldlr^−/−^ll17rc*^Δ*IEC*^) and *ll17rc^flox/flox^* x *Th^cre^* (referred in the text as *ll17rc^ThCre^*) mice. All mice were on C57BL/6 background and were bred in house. *Ldlr^−/−^ll17rc*^Δ*IEC*^ *and Ldlr^−/−^ll17rc^WT^*littermate mice were obtained from the breeding where females were always IL17RC sufficient and did not contain VilCre deleter, therefore excluding generational accumulation and passage of “bad microbiota” from mothers.

Beginning at ∼6 weeks of age, female and male *Ldlr^−/−^ll17rc*^Δ*IEC*^ and *Ldlr^−/−^ll17rc^WT^* littermate mice were placed on Western diet (WD) (Envigo TD 88137: 15.2% kcal from protein, 42.7% kcal from carbohydrate, 42% kcal from fat, 0.2% cholesterol) for 16 weeks. Mice were housed under specific pathogen-free conditions in an AAALAC-approved barrier facility at CSMC. Mice were genotyped by standard PCR protocols.

*To determinate how microbiota can affect atherosclerosis development Ldlr^−/−^ll17rc*^Δ*IEC*^ and *Ldlr^−/−^ll17rc^WT^* littermate mice were separately housed according to genotype or co-housed after weaning.

*Antibiotic treatment experiment.* At 12 weeks of WD feeding *Ldlr^−/−^ll17rc*^Δ*IEC*^ and *Ldlr^−/−^ll17rc^WT^* mice were provided with water containing a mix of broad-spectrum antibiotics: ciprofloxacin 0.2g/L, neomycin 1g/L, vancomycin 0.5g/L, ampicillin 1g/L, metronidazole 0.5g/L, primaxin 0.5g/L.

*In vivo antibody administration.* At 12 weeks of WD feeding *Ldlr^−/−^ll17rc*^Δ*IEC*^ and *Ldlr^−/−^ll17rc^WT^*mice were injected intraperitoneally (i.p.) every 3 days for 4 weeks with mouse anti-TCRγδ antibody (UC7-13D5, Bio X-Cell) (400μg/mouse per injection) or IgG1 isotype control antibody in 3 independent experiments.

*6-hydroxydopamine administration.* At 12 weeks of WD feeding *Ldlr^−/−^ll17rc*^Δ*IEC*^ and *Ldlr^−/−^ Il17rc^WT^* mice were injected intraperitoneally (i.p.) with 6-OHDA 100 mg/kg at day 0 and 150 mg/kg at day 2 or PBS, followed by weekly injections with 150 mg/kg.

*Adeno Associated Virus expressing CRISPR targeting the LDL Receptor in vivo.* AAV serotype 2/8 was produced as previously reported^31^ to generate *Ldlr* AAV-CRISPR from Addgene plasmid #206860. 8 weeks-old mice (*ll17rc^Thcre^* and *ll17rc^WT^*) were injected intraperitoneally (i.p.) with 5×10^11^ genome copies of an Adeno Associated Virus expressing CRISPR targeting the LDL Receptor.

Knock-out of LDL Receptor was confirmed by qPCR on liver upon tissue collection and assessment of serum cholesterol level after 16 weeks of WD feeding.

*Microbiota transfer experiments.* Microbiota recipient *ll17rc^Thcre^*and *ll17rc^WT^* mice were maintained for 8 days with a mix of broad-spectrum antibiotics in drinking water (ciprofloxacin 0.2g/L, neomycin 1g/L, vancomycin 0.5g/L, ampicillin 1g/L, metronidazole 0.5g/L and primaxin 0.5g/L). 24 hours before microbiota transfer, recipient mice were placed on regular water. Microbiota from cecum of *Ldlr^−/−^ll17rc^WT^* or *Ldlr^−/−^ll17rc*^Δ*IEC*^ mice with developed atherosclerosis were weighted, resuspended at concentration 1g of feces/50ml of PBS and gavaged in PBS in 3 consecutive days to *ll17rc^Thcre^* or *ll17rc^ThWT^* mice followed by WD feeding for 16 weeks.

### Quantification of aortic atherosclerotic lesions

Aortic roots were isolated, frozen in an Optimal Cutting Temperature (O.C.T.) compound (Tissue Tek (Sakura)) on dry ice and stored at -80°C. Five-micrometer frozen tissue sections were cut starting at the aortic valve plane to cover 300 μm in intervals of 50μm and stained with Oil Red O/Hematoxylin/Light green as previously described. Images were obtained using Evos FL2 microscope with a 4×0.2 NA objective. The lesion area was measured using Fiji software; atherosclerosis lesion area was calculated as average of lesion areas at least 8 sections for each mouse.

### Immune cells composition analysis by Flow Cytometry

Aortas were isolated from *Ldlr^−/−^ll17rc*^Δ*IEC*^ and *Ldlr^−/−^ll17rc^WT^* mice, cut into small pieces followed by digesting in 2 ml of enzymatic cocktail, containing 450 U/ml collagenase type I, 250 U/ml collagenase type XI, 120 U/ml hyaluronidase type I, 120 U/ml DNAse I in 1x HBSS and incubated in a shaker at 37°C for 55 min. Obtained single cells suspension was stained with following antibodies: CD45-PerCP (30-F11), CD11b-eFluor 450 (M1/70), CD11c-Cy7PE (N418), TCRβ-eFluor 780 (H57–597), CD4-PE (GK1.5), CD8-BV605 (53-6.7), TCRɣδ-FITC (GL3), IL17a-Cy7PE (TC11-18H10.1), IL6-PE (MP5-20F3) and LIVE/DEAD UV fixable dye (Biolegend, 423108) and analyzed by flow cytometry (Symphony A5, BD Biosciences).

Lamina propria lymphocyte (LPL) and Intestinal epithelial lymphocyte (IEL) isolation was performed as described before^64^. In brief, isolated intestines were removed and placed in ice-cold 5% fetal bovine serum (FBS) in HBSS and incubated twice in 20ml of 5mM EDTA in HBSS for 15 min at 37°C with rotation of 150 rpm. After incubation, IELs were collected by passing through a metal sieve into new tubes and the residual tissues (LPL) were chopped to make it almost homogeneous and digested in 5–6 ml collagenase type VIII (Sigma) (1mg/ml) in 5% FBS, HBSS for 35 min in shaking 37°C incubator. After incubation the solution was passed through 70μm cell strainer. Percoll gradient (40%/80%) separation of LPL and IEL was performed by centrifugation at 2200rpm for 25 min with slow acceleration and deceleration. LPL and IEL immune cells were collected from the interphase of Percoll gradient. Obtained cells suspension was stained with following antibodies: CD45-PerCP (30-F11), CD11b-eFluor 450 (M1/70), TCRβ-eFluor 700 (H57–597), CD4-APC (GK1.5), CD8-BV605 (53-6.7), TCRɣδ-FITC (GL3), IL17A-Cy7PE (TC11-18H10.1) and LIVE/DEAD UV fixable dye (Biolegend, 423108) and analyzed by flow cytometry (Symphony A5, BD Biosciences).

For T-cell related cytokine production analysis, single-cell suspensions from aorta were stimulated in complete RPMI 1640 media with PMA + Ionomycin and Brefeldin A for 5h in vitro at 37°C in a CO_2_ incubator. For myeloid cells cytokines staining, cells were stimulated in complete DMEM media with LPS (100ng/ml) for 4h in the presence of Brefeldin A. After stimulation cells were fixed and permeabilized by Cytofix/Cytoperm kit (BD Biosciences) and stained with IL-17A-Cy7PE (eBio17B7), IFNγ-APC (XMG1.2) or IL6 antibody. Flow cytometry data were analyzed using FlowJo software.

### Gene expression analysis

The tissues (aorta, intestine or spleen) were homogenized with RNAse/DNase free 2.8mm Ceramic Beads using Omni Bead Ruptor 24 in PureZOL RNA Isolation Reagent (Bio-Rad Laboratories) followed by RNA isolation using Aurum Total RNA Fatty and Fibrous Tissue kit (Bio-Rad Laboratories) according to manufacturer’s protocol.

First strand cDNA was synthetized using the iScript Reverse Transcription Supermix (Bio-Rad Laboratories). Gene expression was analyzed by SYBR green real-time polymerase chain reaction (Bio-Rad Laboratories) using primers for *Rpl32* (FW 5’-TTCCTGGTCCACAATGTCAA-3’; REV 5’-GGCTTTTCGGTTCTTAGAGGA-3’), *Ccl2* (FW 5’-ATTGGGATCATCTTGCTGGT-3’; REV 5’-CCTGCTGTTCACAGTTGCC-3’), *Il6* (FW 5’-ACCAGAGGAAATTTTCAATAGGC-3’; REV 5’-TGATGCACTTGCAGAAAACA-3’), *Spp1* (FW 5’- TGGCTATAGGATCTGGGTGC- 3’; REV 5’-ATTTGCTTTTGCCTGTTTGG-3’), *Trem2* (FW 5’- CTGGAACCGTCACCATCACTC-3’; REV 5’-CGAAACTCGATGACTCCTCGG-3’), *Il1b* (FW 5’-GGTCAAAGGTTTGGAAGCAG-3’;REV 5’-TGTGAAATGCCACCTTTTGA-3’), *Il17a* (FW 5’-TGAGAGCTGCCCCTTCACTT-3’; REV 5’-ACGCAGGTGCAGCCCA-3’), *Tnf* (FW 5’- AGGGTCTGGGCCATAGAACT-3’; REV 5’-CCACCACGCTCTTCTGTCTAC-3’), *Trgc1* (FW 5’- TCCCCCAAGCCCACTATTTTC-3’; REV 5’- TTCAGCGTATCCCCTTCCTG -3’), *Th* (FW 5’-TGGCTGCCCTCCTCAGTTCT-3’; REV 5’-TGGTCCAGGTCCAGGTCAGG-3’), *Chrb4* (FW 5’-TGGATGATCTCCTGAACAAAACC-3’; REV 5’-CAGGCGGTAGTCAGTCCATTC-3’), *Omg* (FW 5’-TGATAGGGCTGTGGTGGCCT-3’; REV 5’-GCCACGAGCTATGCTGAGGG-3’), *Pirt* (FW 5’-AAGGACGGGAGGCCACAGAT-3’; REV 5’-AACAGGGAGGAGGGAGGCTG- 3’), *Npy*(FW 5’-CAAGAGCAACAACTCGGCATT-3’; REV 5’- GAGAGGGACAGGTTGGCAATC-3’), *Dbh* (FW 5’-GAGGCGGCTTCCATGTACG-3’; REV 5’-TCCAGGGGGATGTGGTAGG-3’), *Gap43* (FW 5’-TGGTGTCAAGCCGGAAGATAA-3’; REV 5’-GCTGGTGCATCACCCTTCT-3’), *Tjp1* (FW 5’-GCGCGGAGAGAGACAAGATG-3’; REV 5’-CTGTGAAGCGTCACTGTGTG-3’), *Tjp2* (FW 5’-GCCTACGAGAAGGTTCTGCT-3’; REV 5’-AGATCCGGCATCTTTGGGTT-3’’), *Ocln* (FW 5’-GTGAGCACCTTGGGATTCCG-3’; REV 5’-TTCAAAAGGCCTCACGGACA-3’), *Cldn2* (FW 5’-CTCTTCGAAAGGACGGCTCC- 3’; REV 5’-CAGTGTCTCTGGCAAGCTGA-3’), *Cldn3*(FW 5’- GACCGTACCGTCACCACTAC-3’; REV 5’-CAGCCTAGCAAGCAGACTGT-3’), *Cldn7* (FW 5’-AATGTACGACTCGGTGCTCG-3’;REV 5’-GTGTGCACTTCATGCCCATC-3’), *Ifnb1* (FW 5’-CCCTATGGAGATGACGGAGA-3’; REV 5’-CCCAGTGCTGGAGAAATTGT-3’), *Cd36* (FW 5’-CAGATCCGAACACAGCGTAG-3’; REV 5’-GCGACATGATTAATGGCACA-3’), *Sra* (FW 5’-ACGACCCGCCACAATTCTC-3’; REV 5’-CTGGAAGCCTTACTTGAAGGAG-3’), *16SrRNA*(FW 5’-ACTCCTACGGGAGGCAGCAGCAGT-3’; REV 5’- TATTACCGCGGCTGCTGGC-3’) (mouseprimerdepot.nci.nih.gov, pga.mgh.harvard.edu/primerbank). Gene expression was normalized to *Rpl32* expression. Data were analyzed using Prism statistical software (GraphPad).

### Bulk RNA sequencing and analysis

Isolated RNA samples were submitted to Novogene Ltd. (Sacramento, USA), where library construction and sequencing were performed. Sequencing was conducted on the Illumina NovaSeq 6000 platform, generating 150 bp paired end reads, with an average of 4 Gb raw data per sample. RNA-seq data was aligned using STAR ^65^ algorithm against mm10 (GRCm38) mouse genome version and RSEM v1.2.12 software ^66^ was used to estimate read counts and FPKM values using gene information from Ensemble transcriptome version GRCm38.89. Raw counts were used to estimate significance of differential expression difference between two experimental groups using DESeq2^67^. Genes ranks proportional to significance were used in GSEA ^68^ analysis. Significantly differentially expressed genes were also analyzed by QIAGEN’s Ingenuity® Pathway Analysis software (IPA, QIAGEN Redwood City,www.qiagen.com/ingenuity) using “Canonical pathways” or “Upstream Regulators” options. FDR<5% results were considered significant unless stated otherwise.

### Mouse single cell RNA sequencing

CD45^+^ cells from aorta or ileum tissue of *Ldlr^−/−^ll17rc*^Δ*IEC*^ and *Ldlr^−/−^ll17rc^WT^*mice fed with WD for 16 weeks were isolated and FACS sorted for CD45^+^Live/Dead^−^ single cells. The single-cell droplets were generated with chromium single-cell controller using Chromium Next GEM Single Cell 3’ Kit v3.1 (10X Genomics, 1000121). Approximately 5,000–10,000 cells were collected to make cDNA at the single-cell level. cDNA was fragmented to ∼270 bp and the Illumina adapters with index, barcode, and UMI were ligated to the fragmented cDNA. After PCR, purification, and size selection, the scRNA libraries were ∼ 450bp and were further sequenced on NovaSeq 6000 (Illumina) at Novogene. Fastq files were obtained for the bioinformatic analysis.

### Bioinformatic analysis of scRNAseq

Using the 10X Genomics Cell Ranger pipeline, reads were aligned to the mouse genome (mm10), and the sequencing depth was subsampled to about 50,000 reads per cell. Further processing, normalization, batch correction, clustering, differential gene expression analysis, and visualization were performed using the Seurat package (v.4.0.5) and the ggplot2 package in R. Contaminant cells were identified and removed through CD45^+^− sorted samples, clustering the cells, and classifying cell type of each cluster. Ileum CD45^+^ cells that were separately sorted and sequenced from LPL and IEL fractions and combined for clustering during bioinformatic analysis. Those cells that have less than 200 or over 5000 unique genes were filtered, and cells that have over 5% mitochondrial counts were filtered. Contaminant CD45 negative cell types were computationally removed from single-cell sequenced CD45^+^ sorted samples. For the combined analysis for aortas cells that have less than 200 or over 4500 unique genes were filtered, and cells that have over 4% mitochondrial counts were filtered. Contaminant CD45^−^negative cell types were computationally removed from single-cell sequenced CD45^+^−sorted samples.

For cell type prediction for cluster classification, the total gene expression of each cluster was analyzed using Gene Set Enrichment Analysis (GSEA) and compared to a gmx file containing gene signatures of all cell types that we derived in house from the Immgen database (www.immgen.org) and T cells and Myeloid cells gene signatures from ^69^. The GSEA analysis was done in the GSEA software^70^. Markers that define clusters (found via differential expression analysis were used to supplement this analysis for cell prediction.

Differentially expressed genes were identified using the FindAllMarkers function from the Seurat R package (v4.0.5). This function was applied to the normalized single-cell expression data to identify cluster-specific marker genes. Parameters were set as follows: only.pos = TRUE to report only genes upregulated in each cluster, min.pct = 0.25 to include genes expressed in at least 25% to filter for genes with a minimum log fold-change of 0.25. The resulting marker gene lists were used for downstream functional enrichment analyses. Statistical comparisons were performed using Fisher’s exact test. These tests were conducted using base R functions (fisher.test) to evaluate whether the distribution of cell types significantly differed between conditions.

### Bioinformatic analysis of human scRNAseq

Publicly available single-cell RNA sequencing data (GSE252243) from patients with coronary artery atherosclerosis were downloaded from the Gene Expression Omnibus (GEO) database. The dataset was processed and analyzed in R using the Seurat package (v4.0.5). Following quality control and normalization, cells from Sample 4 were clustered and visualized using uniform manifold approximation and projection (UMAP). All T cells (CD45 and CD3E positive cells) were extracted were extracted and re-clustered. identified by positive. γδTCR cells specific expression of CD3E, TCRG2, TCRG1. Gene expression levels of IL17A, TCRGC2, and GPR65 were assessed across identified cell clusters using the DotPlot functions.

### Xenium Prime in Situ Gene expression experiment

Fresh frozen aortic root tissue samples from separately housed *Ldlr^−/−^ll17rc^WT^, Ldlr^−/−^ll17rc*^Δ*IEC*^ mice fed with WD for 16 weeks were cut to 6 μm thick sections and profiled using in situ Xenium platform (10x Genomics). The predesigned 5k-gene Xenium Prime 5K Mouse Pan Tissue & Pathways Panel was profiled across the 6 sections by the In Situ Sequencing Infrastructure Unit (10x Genomics), where probe hybridization, ligation and rolling circle amplification were performed according to the manufacturer’s protocol (CG000760 Rev A, 10x Genomics). Background fluorescence was chemically quenched. Imaging and signal decoding were done using the Xenium Analyzer instrument (10x Genomics).

### Bioinformatic analysis of spatial transcriptomics data

Scanpy ^71^ (1.10.3), simpleITK ^72^ (2.3.1), scipy ^73^ (1.12.0) and scvi-tools ^74^(1.2.0) were used to preprocess the raw Xenium spatial transcriptomics data. H&E images of post-Xenium slides were annotated with folded tissue regions and aligned to the spatial transcriptomics data using the Xenium output DAPI image. Data was subsequently preprocessed using the standard Scanpy (1.10.3) pipeline, where cells with median total counts below 20, and genes present in less than 3 unique cells were filtered out. Likewise, Scipy.spatial^73^ (1.2.0) was utilized to remove cells disassociated from the sample via KDTree to exclude all cells with less than 5 cells within a 100um. Expression data was then count normalized to 1000 and natural log transformed. Cell typing was conducted using scvi-tools (1.2.0) and scanpy (1.10.3). Latent spaces were derived from the top 500 highly variable genes in the Xenium scRNA-seq raw gene counts, determined using scanpy.pp.highly_variable_genes (flavor Seurat_v3). The model was formulated using 10 latent variables and 1 hidden layer and was allowed to train for at most 500 epochs (early_stopping = True). Scanpy (1.10.3) was subsequently used to derive a KNN graph of cells from the scVI latent spaces, followed by UMAP embedding, and ultimately leiden clustering (resolution 0.65). Leiden clusters were then assigned cell type labels based on marker gene expression and spatial localization. This process was repeated individually for leiden clusters 0,3, and 11 to further improve immune subtype identification, with the only procedural alteration being the usage of the full 5006 genes rather than just the 500 most variable in scVI latent space generation.

Spatial enrichment of cell type pair (ex. cell type a,b) localization was quantified by computing the spatial cross-correlation as shown in the below equation, using Scipy (1.12.0). All spatial cross-correlations were evaluated at a radius of 100um and used to evaluate the relative spatial enrichment of cell type b in vicinity of cell type a SC(r)= (Observed(r))/(avg(Permuted(r)))-1.

Spatial correlations between cell types a and b were evaluated by building a KDTree centered around a cell type a, followed by counting of all cell type b within a circle of radius r. These were computed for the observed spatial architecture of the tissue, as well as for 100 versions of the data set with unchanged x,y coordinates but randomly permuted cell type labels. The observed value was then divided by the averaged permuted value and subtracted by 1 to determine the over-density (SC(r)> 0) or under-density (SC(r)<0) of cell type b in the proximity of cell type a. All spatial correlations were calculated per sample.

### Immunohistochemistry

For immunohistochemistry staining 7μm thick sections of human atheroma were deparaffinized by taking them through 4 changes in xylene, then washed by 4 changes in 100% ethanol followed by re-hydration in tap water. Antigen retrieval was performed in 1X Citrate buffer (Electron Microscopy Sciences, 64142–08) at 95°C for 1 h followed by 1 h cooling down to RT. Then slides were rinsed in tap water for 3 min and dehydrated in 100% ethanol for 1 min, followed by blocking in 3% H2O2 in PBS for 10 min. Slides were blocked with 5% goat serum in 1% BSA-PBS for 20 min, then they were incubated with primary antibody for TCRγδ (1:30; Fisher Scientific, ENTCR1061), TH (1:50; Sigma, PIMA532984), O/N at 4°C. After washing with 1% BSA-PBS slides were incubated with secondary goat anti-mouse biotinylated antibody (1:125, Biolegend, 405303) for 40 min at RT followed by 40 min of incubation with streptavidin–HRP (1:500; BD Pharmingen, 554066). DAB substrate (Abcam) was applied for 3 min followed by washing in water and counterstaining with Hematoxylin solution (Sigma-Aldrich, HHS32). Excess of Hematoxylin was removed by immersing slides in 0.25% ammonia water followed by rinsing in water. Slides were mounted with coverslips using Permount mounting medium solution (Fisher Chemical, SP15). All images were acquired with EVOS Auto FL2. Microsoft PowerPoint was used for one step brightness adjustment for all images in parallel.

### Immunofluorescent staining analysis

Frozen 7 μm aortic root sections were fixed for 10 minutes in chilled acetone and a solution of 3 mL periodate, lysine, paraformaldehyde (PLP) solution and 1 mL 4% paraformaldehyde was pipetted onto each sample on ice. Sections were washed with phosphate buffered saline (PBS) and incubated with Avidin D/Biotin blocking kit (Vector laboratories) followed by incubation with 5% normal goat serum in 1% BSA in PBS. The sections were stained overnight in 4°C with primary antibodies by co-staining anti-CD11b-FITC (rat, M1/70), anti-CD3 (rat, 17A2) or anti-γδT (rat, GL3) with anti-Th (rabbit, Sigma, AB152), anti-NF200 (sigma,N4142), anti-NPY (Cell Signaling, 11976S) followed by staining with fluorescent conjugated secondary antibodies for 1 hour at room temperature. For co-staining CD11b with Th1, secondary antibodies anti-rabbit AF568 (A10042) (1:400) and anti-FITC AF488 (A11096) (1:1200) were used. For co-staining CD3 with Th1, secondary antibodies anti-rabbit AF488 (A-21206) (1:400) and anti-rat AF568 (A78946) (1:750) were used. Sections were counterstained with DAPI (1:1000) and embedded in Prolong Gold mounting medium. Images were taken using Leica SP8 microscope and analyzed using ImageJ software. Ileum tissues without contents were fixed in Carnoy’s fixative (60% ethanol, 30% chloroform, 10% glacial acetic acid) overnight, paraffin-embedded and stained for Mucin2 with rabbit anti-Muc2 (Santa Cruz, F2) followed by goat anti-rabbit Alexa 488 (Life Technologies) secondary antibody staining.

### PAS staining

PAS staining was performed using a commercial PAS staining kit (Sigma-Aldrich, 1016460001) following the manufacturer’s protocol. Briefly, tissue sections were deparaffinized in xylene, rehydrated through descending grades of ethanol, and rinsed in distilled water. Slides were then incubated in 0.5% periodic acid solution for 10 min at room temperature to oxidize glycols to aldehydes. After rinsing in distilled water, sections were treated with Schiff’s reagent for 15 min in the dark. Following another rinse in distilled water, slides were washed under running tap water for 5–10 min until pink coloration developed. Sections were counterstained with hematoxylin for 1 min, rinsed in tap water, dehydrated through graded ethanol, cleared in xylene, and mounted using a resinous mounting medium. Stained sections were examined under a bright-field microscope (Evos Auto FL2) at various magnifications. PAS-positive structures, such as glycogen, mucopolysaccharides, and basement membranes, were identified by their characteristic magenta coloration. Images were captured using a digital imaging system under consistent exposure settings.

### Tissue Clearing

For whole-mount imaging, the iDISCO aorta clearing method was used with modifications (Renier et al., 2014). Samples were gradually fixed in acetone/H_2_O series in 25% for 1h, 50%, for 3h, 25% 1h, 80%, then washed in PBS with shaking for 30 min 2 times, then PBS/30% sucrose for 2h. After performed decolorization in PBS/30% Sucrose with 1% H_2_O_2_ and 10mM EDTA-Na (pH 8.0) at 4°C overnight. Washed in PBS with shaking for 30 min 2 times and pretreated with PBS with 0.2%Triton X-100, 1% deoxycholate, 10% DMSO, 10mM EDTA overnight. After permeabilization, samples were incubated in PBS with 0.2% Triton X-100, 10% DMSO, and 5% normal donkey serum for 2 days at RT to block tissue. Samples were then incubated with primary antibodies in PBS with 0.1% Tween 20 and 10 mg/mL heparin (PTwH) with 5% normal donkey serum at RT for 48h followed by 5 washes of PTwH, 1 hour each at RT, plus 1 overnight wash at RT. Samples were then incubated with secondary antibodies in PTwH at RT for 48h, followed by 5 washes of PTwH, 1 hour each at RT, plus 1 overnight wash at RT. Tissue clearing was performed after embedding tissue in 0.8% agarose in PBS. Blocks were incubated in methanol/H2O series 20% 1h 3 times, 40%, 60% 80% and 100% 1h each, then 100% overnight. Then samples were incubated with shaking in 66% DCM/33% Methanol at RT and then 100% DCM 30 min 3 times. After samples were incubated in Dibenzyl ether (DBE) 12h 2 times without shaking.

### Light Sheet microscopy Imaging

Whole mount cleared samples were imaged using a LaVision BioTec Light Sheet UltraMicroscope II (Miltenyi Biotec). Samples were immersed in the imaging chamber filled with dibenzyl ether (DBE, RI ≈ 1.56) and mounted on adapter to prevent movement during acquisition. Images were acquired using a MV PLAPO 2x objective lens at 0.63x magnification. Illumination was achieved using a 488 nm laser line to excite fluorescent signals, and detection was performed with an Olympus MVX10 macro zoom system coupled to an Andor Neo cCMOS camera. Z-stack images were collected at 3.9 µm step intervals across the sample volume. All image stacks were post-processed using Imaris (Bitplane) and Fiji (ImageJ) for 3D reconstruction, visualization, and analysis. Laser power and exposure times were optimized for each sample to minimize photobleaching and maximize signal-to-noise ratio.

### Norepinephrine, Acetylcholine and NPY measurement

Norepinephrine, Acetylcholine and NPY levels in aortas of WD-fed *Ldlr^−/−^ll17rc^WT^, Ldlr^−/−^ Il17rc*^Δ*IEC*^ separately housed mice were assessed in aortic supernatants obtained when freshly isolated aortas were incubated on ice for 4h in RPMI-1640 media. Norepinephrine and Acetylcholine levels were assessed in serum of WD-fed *Ldlr^−/−^ll17rc^WT^, Ldlr^−/−^ll17rc*^Δ*IEC*^ separately housed mice or after treatment with a mix of broad-spectrum antibiotics. Blood samples were drawn from mice and serums were collected via centrifugation for 10 min. Serum and aortas supernatant Norepinephrine, Acetylcholine and NPY level was analyzed using Norepinephrine ELISA (Abcam, ab287789), Acetylcholine ELISA (Abcam, ab287812) and NPY ELISA (Sigma, EZRMNPY-27K) according to manufacture protocol and analyzed using spectrophotometer (CLARIOstar plate reader, USA).

### NDG and DRG neurons differentiation and treatment

Mix of Nodose ganglions and dorsal root ganglia were seeded into flat-bottom 96-well plates coated with laminin (Thermo Fisher) and allowed to attach to the bottom of the wells for 2 hours, then medium was replaced with neurobasal medium containing 50 ng/μl nerve growth factor (Thermo Fisher). Following an overnight incubation, the supernatant was removed and replaced with neurobasal medium containing 50 ng/μl nerve growth factor, 10 mM cytosine arabinoside (Sigma), and 100 ng/ml IL-17A. Culture medium was replaced daily for the duration of the experiment. At the endpoint, 72h later, the supernatant was removed, cells were lysed with RLT Plus buffer (Qiagen) and the RNA was extracted to perform bulk RNA-seq.

### FITC-dextran permeability test

Intestinal permeability was assessed by FITC-dextran translocation. FITC-dextran (Sigma, 46944). Mice were fasted for 4 h prior to oral gavage. FITC-dextran was administered by gavage at a dose of 600 mg/kg body weight, diluted in phosphate-buffered saline (PBS). After 4 h, blood was collected via retro-orbital bleed, and plasma was isolated by centrifugation. Quantification was performed by comparing FITC fluorescence in the receiving compartment relative to input controls. Background fluorescence from blank wells or untreated tissues was subtracted, and data were normalized to control conditions.

### Metabolomics

Blood was collected from mice at the time of sacrifice, and serum was isolated. Untargeted metabolomic analysis was performed using a gas chromatography time-of-flight mass spectrometer (GCTOF-D) at the UC Davis West Coast Metabolomics Center (WCMC). Metabolome detection intensity levels were quantile normalized and log2-transformed. Significance of changes were tested using Limma package version 3.36.1 ^75^ .

### Serum Lipid Analyses

Blood samples were collected, and serum was isolated by centrifugation at 5000 rcf for 10 minutes. Total cholesterol was determined using the Cholesterol E Assay Kit (Fujifilm, NC9138103) according to manufacture protocol. Absorbance was measured at 600 nm using a spectrophotometer. Triglycerides were determined using the Triglyceride Colorimetric Assay Kit (Cayman Chemical, 10010303) according to manufacture protocol. Absorbance was measured at 550 nm. Low-density lipoprotein (LDL) was determined using the Mouse LDL-Cholesterol Kit (Crystal Chem, 79980) according to manufacture protocol. Absorbance was measured at 600 nm.

### Microbiome analysis

For whole genome shotgun (WGS) sequencing of gut microbiota, bacterial DNA was isolated from a fecal microbiome (ceacum part) of separately housed and co-housed *Ldlr^−/−^ll17rc^WT^* and *Ldlr^−/−^ Il17rc*^Δ*IEC*^ mice. The stools were homogenized with RNAse/DNase free 2.8mm Ceramic Beads using Omni Bead Ruptor 24 in ASL buffer (stool lysis buffer) (Qiagen, 51604) followed by DNA isolation using QIAamp DNA stool Mini Kit (Qiagen, 51604) according to manufacturer’s protocol. Quality control, library construction, and sequencing procedures were carried out at Nogogene according to Novogene’s standard protocols.

### Metagenomic whole genome shotgun (WGS) sequencing analysis

The analysis of the shotgun sequencing data was done as described before^76^. Metagenomic samples were processed using the JAMS package version 1.9.8 with R/4.4.1, available at https://github.com/johnmcculloch/JAMS_BW^77^ using the JAMSdb202212 database. Shorthands for filtering presets are available with “light” filtering meaning that only LKTs with relative abundances of more than 50 PPM in 5% of samples, taxonomy information must come from >70% contigs in at least 50% of samples AND an estimated genome completeness of 5% within 5% of the samples within the SummarizedExperiment object are kept. All other taxonomic features (LKTs) not fulfilling these criteria were discarded when this filter is applied.

### 16S RNA sequencing

To analyze 16S rRNA of the adhesive bacteria from different parts of intestine, RNA was isolated from separately housed and co-housed *Ldlr^−/−^ll17rc^WT^* and *Ldlr^−/−^ll17rc*^Δ*IEC*^ mice by RNA isolation using Aurum Total RNA Fatty and Fibrous Tissue kit (Bio-Rad Laboratories) according to manufacturer’s protocol. cDNA synthesis and 16S V4 rRNA gene sequencing were performed by Novogene Co., Ltd. (Sacramento, USA). Quality control, library construction, and sequencing procedures were carried out at Nogogene according to Novogene’s standard protocols. Paired-end fastq reads were subsequently analyzed using DADA2 ^78^ and QIIME 2 2019.4 ^79^.

Batch corrections for cohorts (different experiments) were performed using ComBat in the sva package. For comparisons between samples, ordination plots were made with the t-SNE algorithm using the uwot package in R (https://github.com/jlmelville/uwot) and the ggplot2 library. PERMANOVA values were obtained using the adonis function of the vegan package, with default (999) permutations and pairwise distances calculated using Bray–Curtis distance. Heat maps were drawn using the ComplexHeatmap package in R. For each feature, P values were calculated using the Mann–Whitney–Wilcoxon U test on ppm relative abundances for that feature in samples within each group. The log2FC values shown on the heat maps were obtained by calculating the fold change between the geometric mean ppm relative abundance for samples within each group, using **Statistical analysis**

Student’s two-tailed t-test was used for comparison between two groups, ANOVA test was used for multiple comparisons. Data were analyzed using the GraphPad Prism Software (Version 7.0). Data are presented as mean ± SEM; *p< 0.05, **p < 0.01, ***p < 0.001, ****p<0.0001. A p-value < 0.05 was considered statistically significant.

### Data Availability

The data generated in this study are available upon reasonable request from the corresponding author. RNAseq data generated in this study were deposited to Gene Expression Omnibus (GEO) with an accession number GSE322757.

## Supplemental Figures

**Figure S1.**
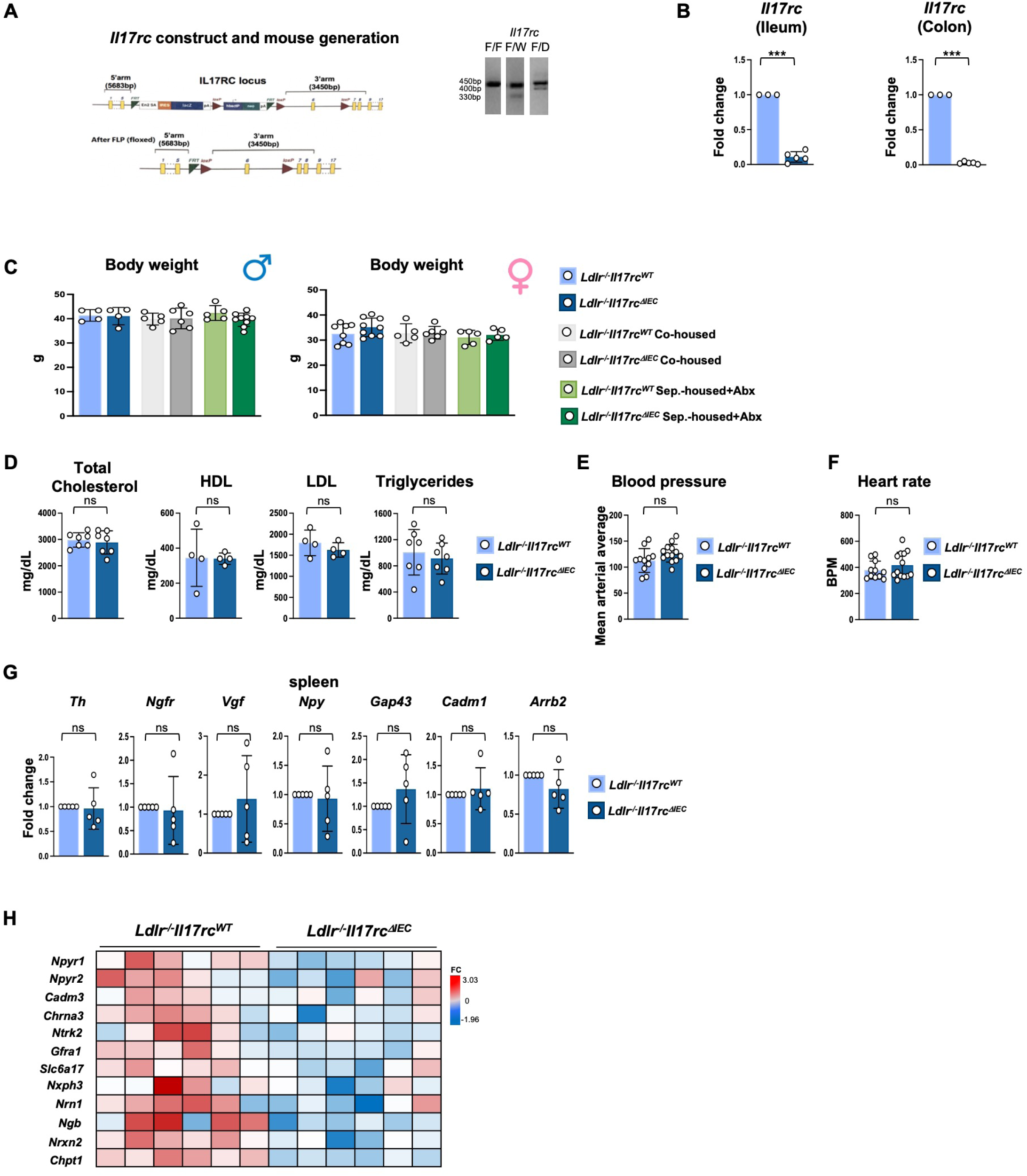
Loss of intestinal epithelial cell Il-17RC promotes atherosclerosis but does not affect body weight, blood pressure and serum lipid profile. Related to Figure 1. **A.** Scheme of IL17RC targeting construct to generate *ll17rc^fl/fl^* mice along with genotyping results of *ll17rc^fl/fl^, ll7rc^fl/w^, ll7r^fl/d^, ll17rc^w/w^* alleles. **B.** Q-RT-PCR analysis of *ll17rc* gene expression in Ileum and colon from *Ldlr^−/−^ ll17rc^WT^* and *Ldlr^−/−^ ll17rc^ΔIEC^* mice. Gene expression was normalized to *Rpl32* and then to gene expression in *Ldlr^−/−^ll17rc^WT^.* Data are mean ± SEM from 3 independent experiments. ***p<0.001, Student’s t-test. **C.** Body weight of *Ldlr*^−/−^*ll17rc^WT^* and *Ldlr*^−/−^*ll17rc^ΔIEC^* separately housed, co-housed and +ABX mice after 16 weeks on WD (n=5-10), two-way ANOVA followed by Tukey’s post-hoc test was applied. **D.** Serum triglyceride, high-density lipoprotein (HDL), low-density lipoprotein (LDL) and total cholesterol levels in *Ldlr^−/−^ll17rc^WT^* and *Ldlr*^−/−^*ll17rc^ΔlEC^* separately housed mice. Blood pressure (E) and **(F)** heart rate of *Ldlr*^−/−^*ll17rc^wr^ and Ldlr^−/−^IH7rc^ΔIEC^* separately housed mice after 16 weeks on WD (n=9-12 mice), Student’s t-test. G. Expression of selected neuronal genes in spleen from separately housed *Ldlr*^−/−^*ll17rc^WT^* (n=5) and *Ldlr*^−/−^*ll17rc^ΔIEC^* (n=5) mice. Gene expression was normalized to *Rpl32* and then to gene expression in *Ldlr*^−/−^*ll17rc^WT^.* Data are mean ± SEM from 3 independent experiments. Student’s t-test. I. Bulk RNA sequencing analysis with a heatmap of differentially expressed genes selected neuronal genes in ileum samples from of *Ldlr^−/−^ll17rc^WT^* and *Ldlr*^−/−^*ll17rc^ΔIEC^* mice (n=6). Red indicates upregulated and blue indicates downregulated gene expression levels, p<0.05.FC-Fold change

**Figure S2.**
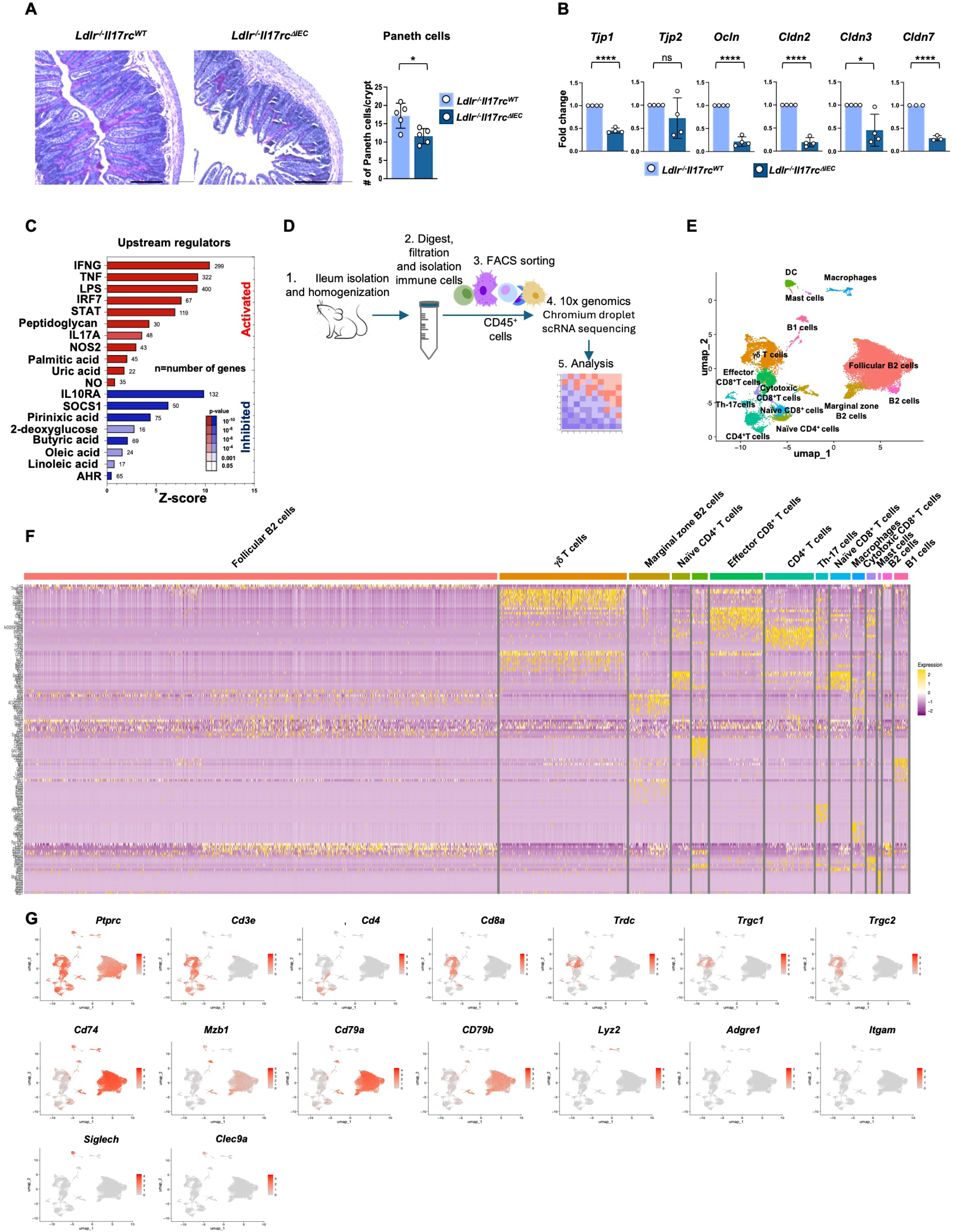
Loss of IL-17RC signaling in intestine reduces barrier function and enhances intestinal inflammation. Related to Figure 2. **A.** Representative images of PAS staining in ileum and quantification of Paneth cells number in separately housed *Ldlr^−/−^ll17rc^WT^* and *Ldlr^−/−^ll17rc^ΔIEC^* mice (n=5) fed by WD for 16 weeks. Scale bar is 50 pm. B. Expression of selected tight junction proteins genes in the ileum (n=4-5) from *Ldlr^−/−^ll17rc^WT^* and *Ldlr^−/−^ll17rc^ΔIEC^* mice. Gene expression was normalized to *Rpl32* and then to gene expression in *Ldlr^−/−^ll17*rc*^WT^ as* determined by Q-RT-PCR. Data are mean ± SEM from 3 independent experiments. *p<0.05, ****p<0.0001, Student’s t-test. **C.** Upregulated (red) and downregulated (blue) upstream regulators as determined by IPA analysis. **D.** Scheme of experiment. Ileum tissue from *Ldlr*^−/−^*ll17rc^WT^* and *Ldlr^−/−^ll17rc^ΔIEC^* mice fed with WD for 16 weeks were dissociated into single cell suspension and FACS sorted for CD45^+^ cells and applied for 10X Genomics Chromium droplet platform for cell isolation and sequencing. Analysis of CD45^+^ cells revealed the presence of 17 clusters comprising 14 different immune cell types. Clustering analysis revealed presence of 6491 T cells among 13299 sorted CD45^+^ cells which we identified 3 clusters of CD4 (1333 cells) and 2 clusters of CD8 cells (1616 cells) and gdT cells (3542). E. UMAP further confirmed the specificity of clusters **F.** Heatmap displays top marker genes defining each cluster. **G.** UMAP plots for cluster-defining genes for CD45^+^ cells.

**Figure S3.**
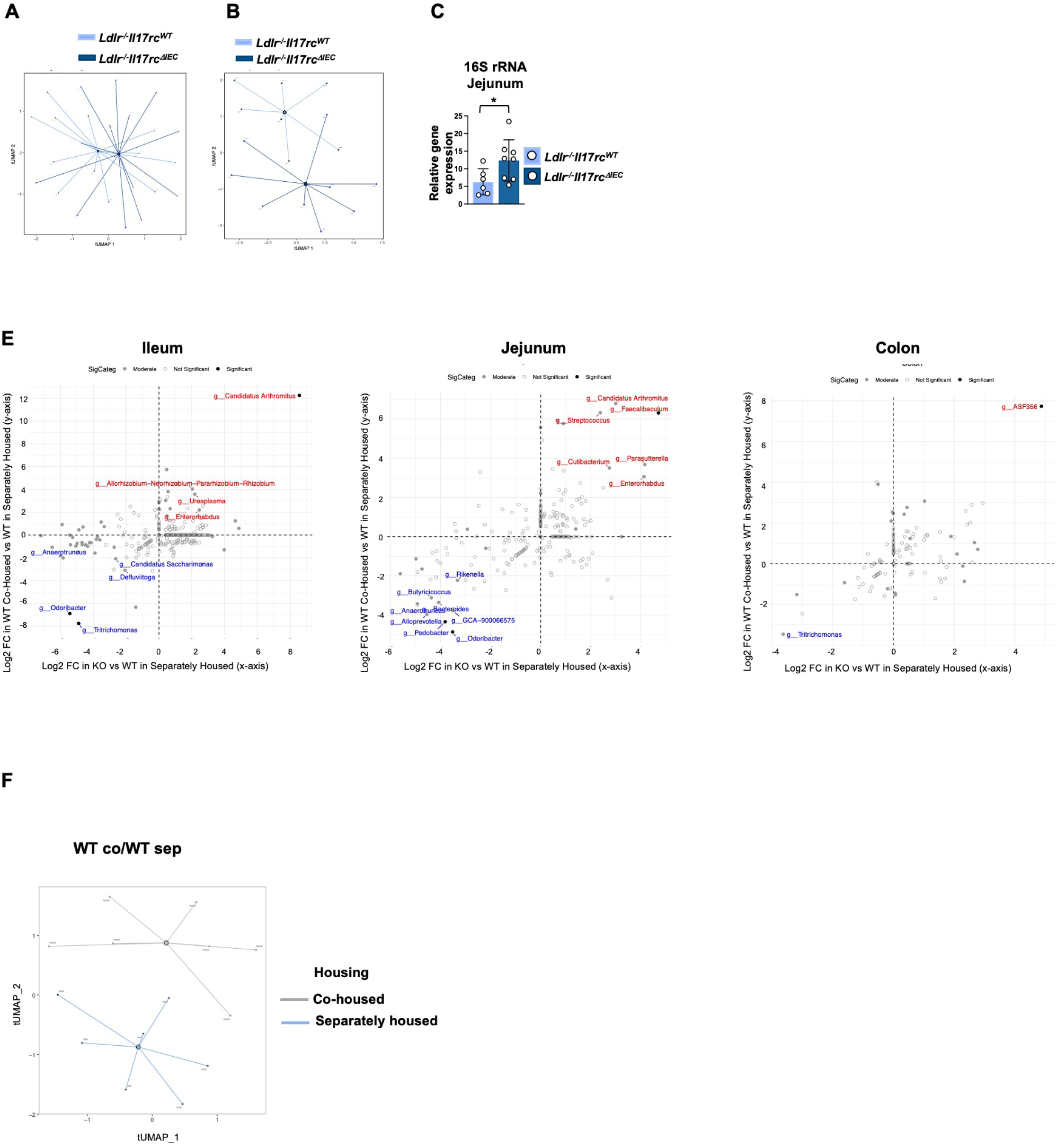
Intestinal IL-17RC regulates luminal and adhesive microbiome composition. Related to Figure 2 and 3. **A.** Differential luminal microbiome composition in *Ldlr^−/−^ll17rc^WT^* and *Ldlr^−/−^ll17rc^ΔIEC^* mice fed with WD for 16 weeks as determined by shotgun metagenomics sequencing followed by Bray-Curtis t-distributed uniform manifold approximation and projection (tUMAP), p<0.0462; **B.** Bray-Curtis t-distributed uniform manifold approximation and projection (tUMAP) of tissue adhesive bacteria in *Ldlr^−/−^ ll17rc^WT^* and *Ldlr^−/−^ll17rc^ΔIEC^* mice after fed with WD for 16 weeks as determined by 16S rRNA sequencing, p<0.0015. **C.** 16S rRNA quantification in jejunum tissue from separately housed *Ldlr*^−/−^ *ll17rc^WT^* and *Ldlr*^−/−^*ll17rc^ΔIEC^* (n=7) mice. Gene expression was normalized to *Rpl32.* Data are mean ± SEM from 3 independent experiments. *p<0.05, Student’s test. **D.** Selected bacteria load from Jejunum and Ileum from *Ldlr^−/−^ll17rc^WT^* and *Ldlr*^−/−^*ll17rc^ΔIEC^* mice (n=5-8) as determined by 16S rRNA sequencing of isolated and washed tissue, p<0.05, Student’s t-test. **E.** Scatter plots comparing adhesive bacteria in ileum, jejunum and colon tissues. X axis indicates Iog2 fold change in *Ldlr*^−/−^*ll17rc^WT^* and *Ldlr^−/−^ll17rc^ΔIEC^* mice after 16 weeks of WD feeding as determined by 16S rRNA sequencing. The Y axis indicates Iog2 fold change in co-housed *Ldlr^−/−^ll17rc^WT^ vs* separately housed *Ldlr*^−/−^*ll17rc^WT^.* Significant (black circle), moderate (Grey circle) and not significant (empty circle) is if the bacteria have p-value < 10% in both, either, or neither comparison, respectively. Red - upregulated bacteria, blue - downregulated bacteria, p<0.05. **F.** Bray-Curtis t-distributed uniform manifold approximation and projection (tUMAP) of tissue adhesive bacteria in *Ldlr*^−/−^*l17rc^WT^ co-housed* with *Ldlr*^−/−^*ll17rc^ΔIEC^* and *Ldlr*^−/−^*ll17rc^WT^* separately housed mice as determined by 16S rRNA sequencing.

**Figure S4.**
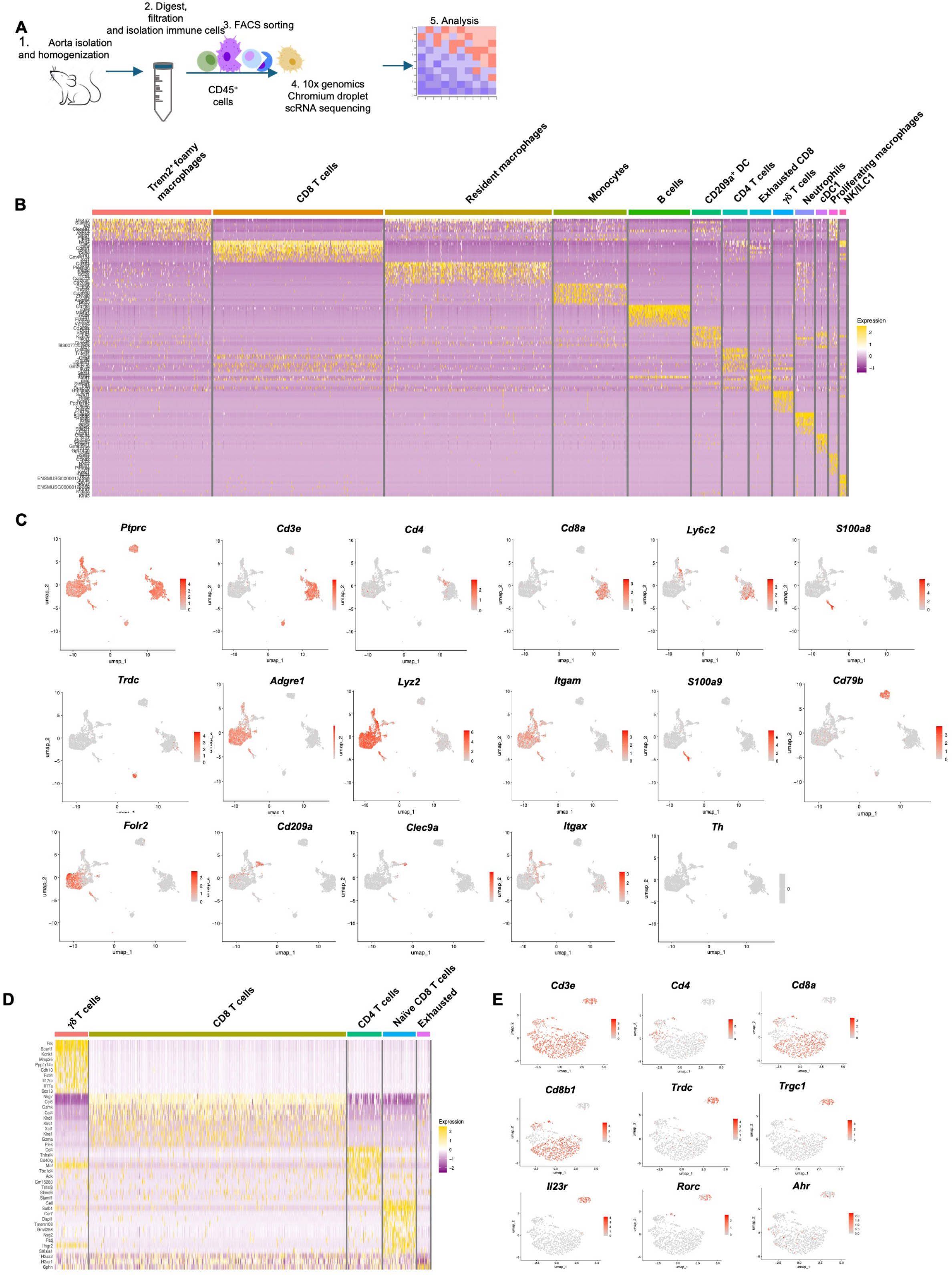
Single cell RNA sequencing analysis of FACS-sorted immune cells from aorta. Related to Figure 4. **A.** Scheme of scRNA seq pipeline to analyze CD45^+^ immune cells in aorta from separately housed *Ldlr^−/−^ll17rc^WT^* and *Ldlr^−/−^ll17^ΔEC^* mice fed with WD for 16 weeks. Aortas were dissociated into single-cell suspensions, and CD45^+^ cells were sorted by FACS and applied to single­cell RNA sequencing using 10x Genomics Chromium droplet platform. Cells clustering revealed presence of 2913 cells among which we identified 13 clusters. B. Heatmap of top differentially expressed cluster-defining marker genes. **C.** UMAP plots for cluster-defining genes among CD45^+^ cells. Analysis of scRNA seq of aortic CD45^+^ cells clusters revealed the presence of 944 T cells among 2913 sorted CD45^+^ cells; among which we identified 2 clusters of CD4 (172 cells) and 2 clusters of CD8 cells (688 cells) and gd T cells (84). Cell type-specific gene expression analysis further confirmed the specificity of clusters. D. Heatmap of top differentially expressed genes defining T cell clusters. **E.** UMAP plots for selected genes for T cells among sorted CD45^+^ cells.

**Figure S5.**
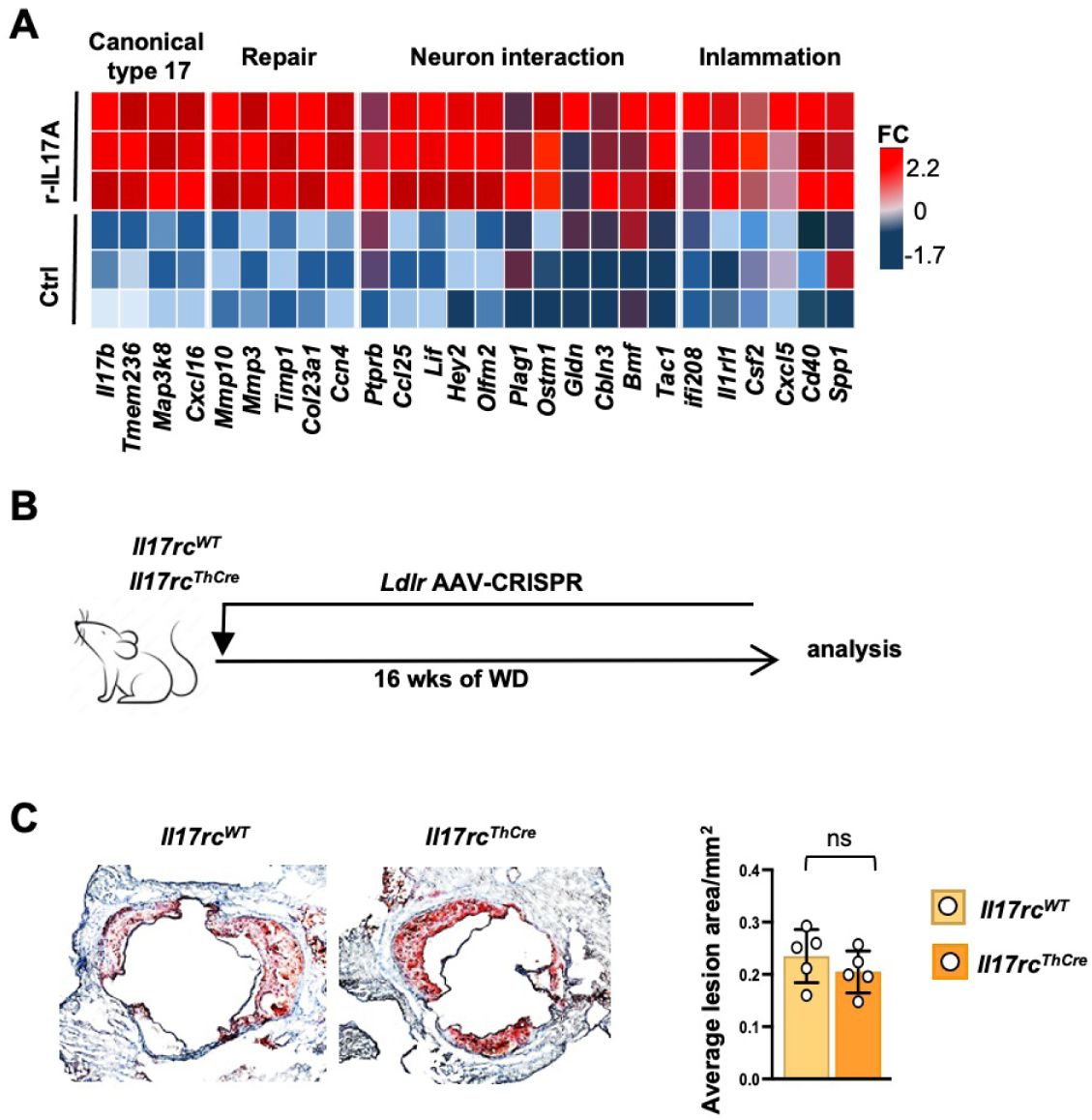
Sympathetic IL17RC deficiency does not affect atherosclerosis in the absence of intestinal perturbations. Related to Figure 5. **A.** Heatmap shows significantly changed genes as determined by bulk RNA sequencing after stimulation of mix of neurons isolated from dorsal root ganglia and nodosal ganglia with r-IL17Afor 72h. FC-Fold change. B. Scheme of experiment. *H17rc^WT^* and *IH7rc^Thcre^* mice were administered with AAV-CRISPR-LDLR and fed with WD for 16 weeks. **C.** Images of aortic arch and representative images of aortic root sections and quantitative comparison of atherosclerotic lesion sizes between *1117rc^m^ and IH7rc^Thcre^* mice (n=5), Student’s t-test.

**Figure S6.**
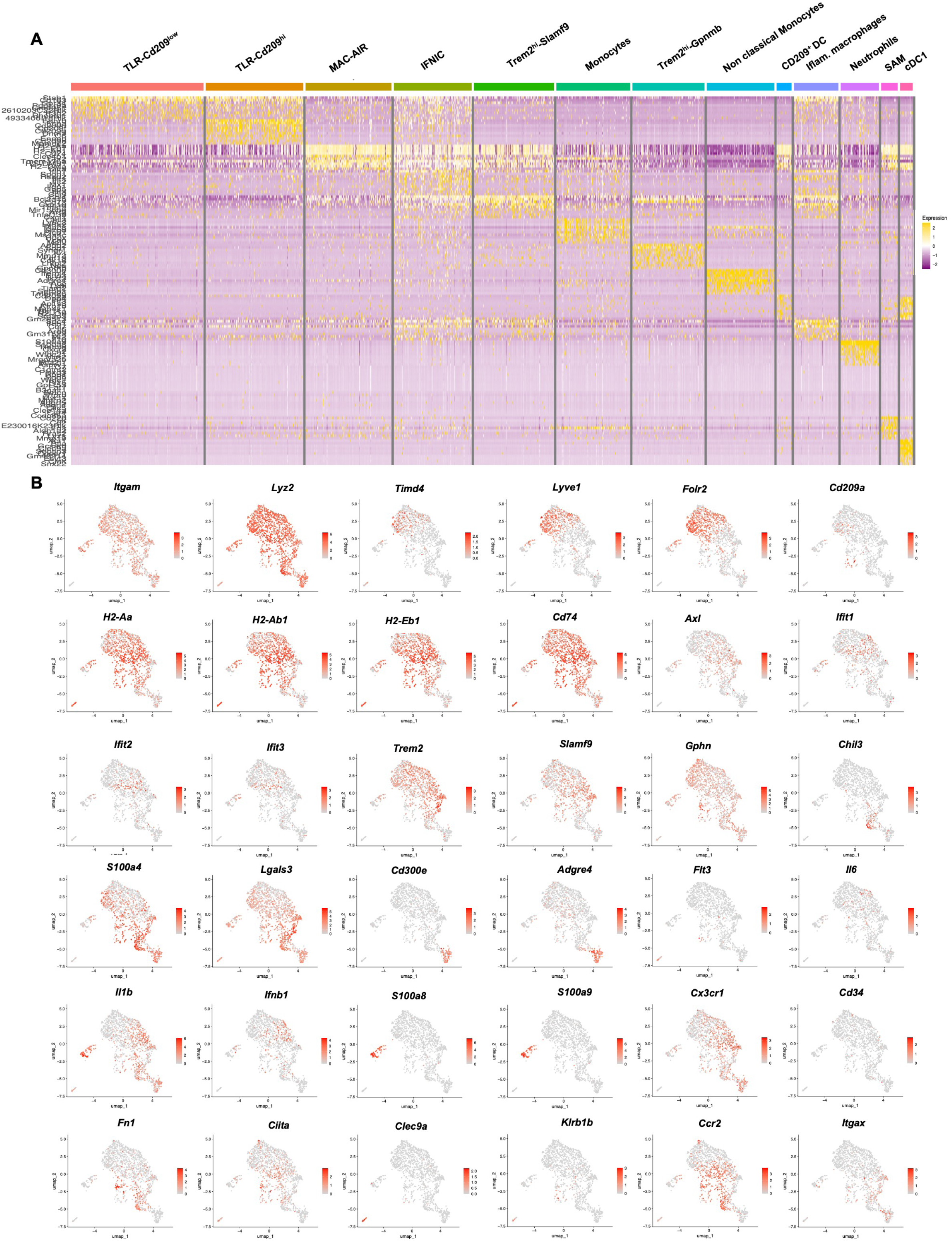
Single cell RNA sequencing analysis of extracted myeloid cells from aorta. Related to Figure 6. Clustering analysis revealed presence of 1411 myeloid cells among 2913 sorted CD45^+^ cells, among which we identified 8 clusters of macrophages (1052 cells), 3 clusters of monocytes (271 cells), 1 cluster of neutrophils (66 cells) and DC cells (22 cells). A. Heatmap of top cluster-defining genes. B. UMAP plots for cluster-defining genes for myeloid cells.

**Figure S7.**
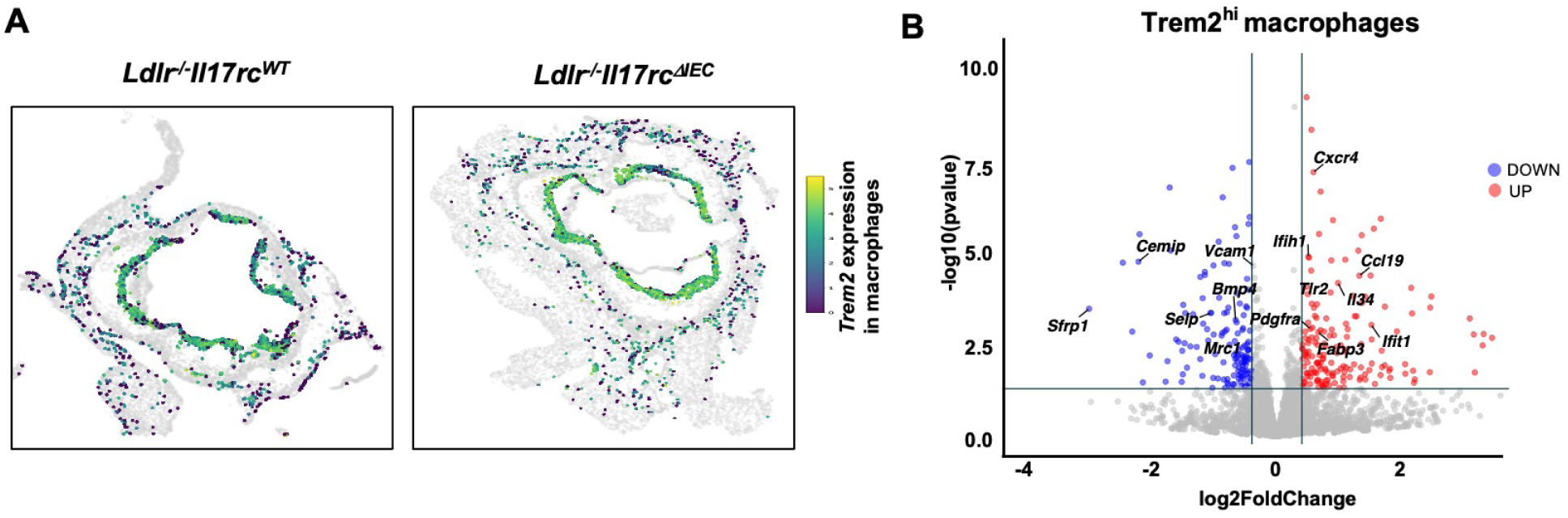
Intestinal IL-17RC regulates macrophage activation in atherosclerosis. Related to Figure 6. **A.** Representative Xenium spatial transcriptomic images of the aortic root from *Ldlr^−/−^ll17rc^WT^* and *Ldlr^−/−^ll17rc^ΔIEC^* mice demonstrate Trem2^+^ macrophage localization. **B.** Volcano plot characterizing gene expression differences in *Trem2^hi^* macrophages between *Ldlr^−/−^ll17rc^ΔIEC^* and *Ldlr*^−/−^*ll17rc^WT^* mice, p<0.05.

## Notes

### Competing Interest Statement

The authors have declared no competing interest.

